# Global patterns of helminths associated with gelatinous zooplankton: a missing link in marine parasite transmission

**DOI:** 10.64898/2026.07.29.741456

**Authors:** Anastasiia Iakovleva, Dror L. Angel, Tamar Guy-Haim

## Abstract

Gelatinous zooplankton are abundant predators and prey in marine food webs, yet their role in helminth (parasitic worms) transmission remains poorly resolved. Here we combine a global synthesis of published records with new morphological and molecular observations from the Mediterranean, Red, Celtic, Baltic and North Seas to assess the ecological role of helminths associated with gelatinous zooplankton. We compiled 431 host–parasite association records from 89 sources, including 23 new records, and show that helminth occurrence and richness in gelatinous hosts are concentrated at temperate latitudes, contrasting with the classical latitudinal diversity gradient. Our sampling revealed markedly higher parasite prevalence, abundance and diversity in the Red Sea than in the Mediterranean, while no helminths were detected in gelatinous zooplankton sampled from the Baltic and North Seas. All helminths recorded were larval stages, indicating that gelatinous zooplankton function as key intermediate hosts in marine helminth life cycles. Molecular analyses identified cestodes, nematodes and digenean trematodes associated with cnidarians, ctenophores, and chaetognaths, including the first record of trematode larvae in a pelagic tunicate. Our findings challenge the view of gelatinous zooplankton as dead-end hosts and identify them as overlooked vectors that may shape marine parasite biogeography, food web connectivity and invasion dynamics.

## Main

Parasites are among the most abundant and evolutionary successful life forms on Earth, occurring in every ecosystem and infecting hosts from all major taxonomic groups^1^. Although traditionally viewed primarily through the lens of disease, parasites are now increasingly recognized as integral components of ecological communities and important drivers of ecosystem processes^2–4^. Approximately half of all known species are estimated to live parasitic lifestyles^5^, with helminths – a collective term encompassing parasitic nematodes, digenean trematodes, cestodes, and acanthocephalans – representing one of the largest and most diverse groups of parasites. Far from being merely consumers of host resources, helminths perform multiple ecological functions^4^. They can act as ecosystem engineers by modifying host traits and living material in ways that increase resource availability for other organisms, contribute to the maintenance of local biodiversity^2,6^, and influence energy flow and food web structure by altering trophic efficiency^7,8^. By exploiting hosts across multiple trophic levels and habitats, helminths enhance connectivity among ecosystems, linking otherwise distinct food webs and facilitating the transfer of energy and matter^9,10^. Although new helminth species descriptions from definitive hosts have been rising in the past 40 years, studies that characterize the larval stages and identify intermediate hosts remain scarce^11^.

A substantial proportion of helminth life cycles involve planktonic organisms as intermediate hosts, highlighting the importance of understanding parasite interactions within pelagic ecosystems. Gelatinous zooplankton constitute a major component of these planktonic communities, yet their role as helminth hosts remains poorly understood. This functional, non-taxonomic assemblage of organisms is characterized by soft, gelatinous bodies and include cnidarians, ctenophores, pelagic tunicates, and chaetognaths^12^. Given their broad trophic interactions, gelatinous zooplankton are expected to play an important role in marine helminth transmission. Nevertheless, records of helminth infections in gelatinous plankton are sparse and geographically fragmented, with most documented cases involving trematodes^13–17^. Despite growing interest in gelatinous zooplankton and their symbiotic associations^18–21^, knowledge of their parasite diversity remains very limited. To date, no study has comprehensively synthesized the diversity and biogeography of helminths associated with gelatinous zooplankton. This knowledge gap likely reflects both conceptual biases, whereby gelatinous organisms have long been considered trophic dead ends^22^, and methodological challenges associated with their sampling and preservation.

Beyond their influence on ecosystem processes^22–26^, abundant populations of gelatinous zooplankton may facilitate parasite transmission and dispersal over large spatial scales. In particular, pelagic hosts can transport infective stages across habitats via ocean currents and vertical migrations^27,28^, potentially increasing connectivity among parasite populations and between otherwise weakly connected food webs. Understanding parasite transmission through gelatinous zooplankton is particularly important in the context of large-scale community shifts and biological invasions. By altering host survival, reproduction, behavior, and trophic interactions, parasites can influence ecological processes across multiple trophic levels and modify invasion outcomes^29–31^. Depending on ecological context, parasites may either facilitate or constrain bioinvasions through mechanisms such as enemy release, spillover, spillback, or dilution effects^29,32,33^. Co-introduced helminths may also strengthen the interactions among invasive species and intensify their impacts on native communities^34,35^.

The Mediterranean and Red Seas provide a valuable natural model system for addressing these questions. The opening of the Suez Canal initiated one of the world’s largest ongoing marine invasions, resulting in extensive biotic exchange between the Red Sea and Mediterranean Sea^36,37^. Simultaneously, rapid warming and tropicalization of the eastern Mediterranean have facilitated the establishment of non-native species and reshaped community structure, including increases in the abundance and bloom frequency of gelatinous zooplankton^38^. Because gelatinous zooplankton can serve as intermediate or paratenic hosts^18,19,39,40^, these changes may alter parasite transmission pathways in the region and create new opportunities for parasite dispersal across biogeographic boundaries. A key unresolved question is therefore how biological invasions and the associated expansion of gelatinous zooplankton populations affect parasite diversity, transmission pathways, and invasion-related processes on a broader scale. Here, we combine a global synthesis of published records with new observations from the Mediterranean and Red Seas to characterize the diversity and biogeography of helminths associated with gelatinous zooplankton and to assess their potential role in parasite-mediated invasion processes.

## Results

### Biogeographical patterns of gelatinous hosts and their parasites

Our systematic literature search for records of helminths associated with gelatinous zooplankton yielded 1094 publications, the earliest dating to 1830^41^. Of these, 1057 were excluded due to irrelevance (non-marine, non-helminth and/or non-gelatinous host). An additional 52 articles were through manual screening. A total of 431 helminth association records in gelatinous zooplankton were extracted from 89 references, including 23 new records generated in the present study (Supplementary Tables S1, S2).

The global distribution of helminths associated with gelatinous zooplankton is presented in figure 1a. The relative frequency of helminth occurrence (Fig. 1b) and species richness (Fig. 1c) were analyzed in 5° latitudinal bins. Most helminth records were concentrated between 20° and 60° in the Northern Hemisphere and between 25° and 45° in the Southern Hemisphere (Fig. 1b). A similar pattern was observed for helminth species richness (Fig. 1c). Both record frequency (%) and species richness peaked at approximately 30°S, accounting for 22% of all records and 37 helminth species. In the Northern Hemisphere, relatively high values occurred between 20° and 60°, with 8–12% of records and 9–18 species per latitudinal bin. In contrast, tropical regions showed relatively few records (0–2%) and low species richness (0–4 species; Fig. 1b, c). Across all records, the parasite assemblage was strongly dominated by trematodes, followed by cestodes, nematodes, and acanthocephalans (Fig. 1d).

**Figure 1.**
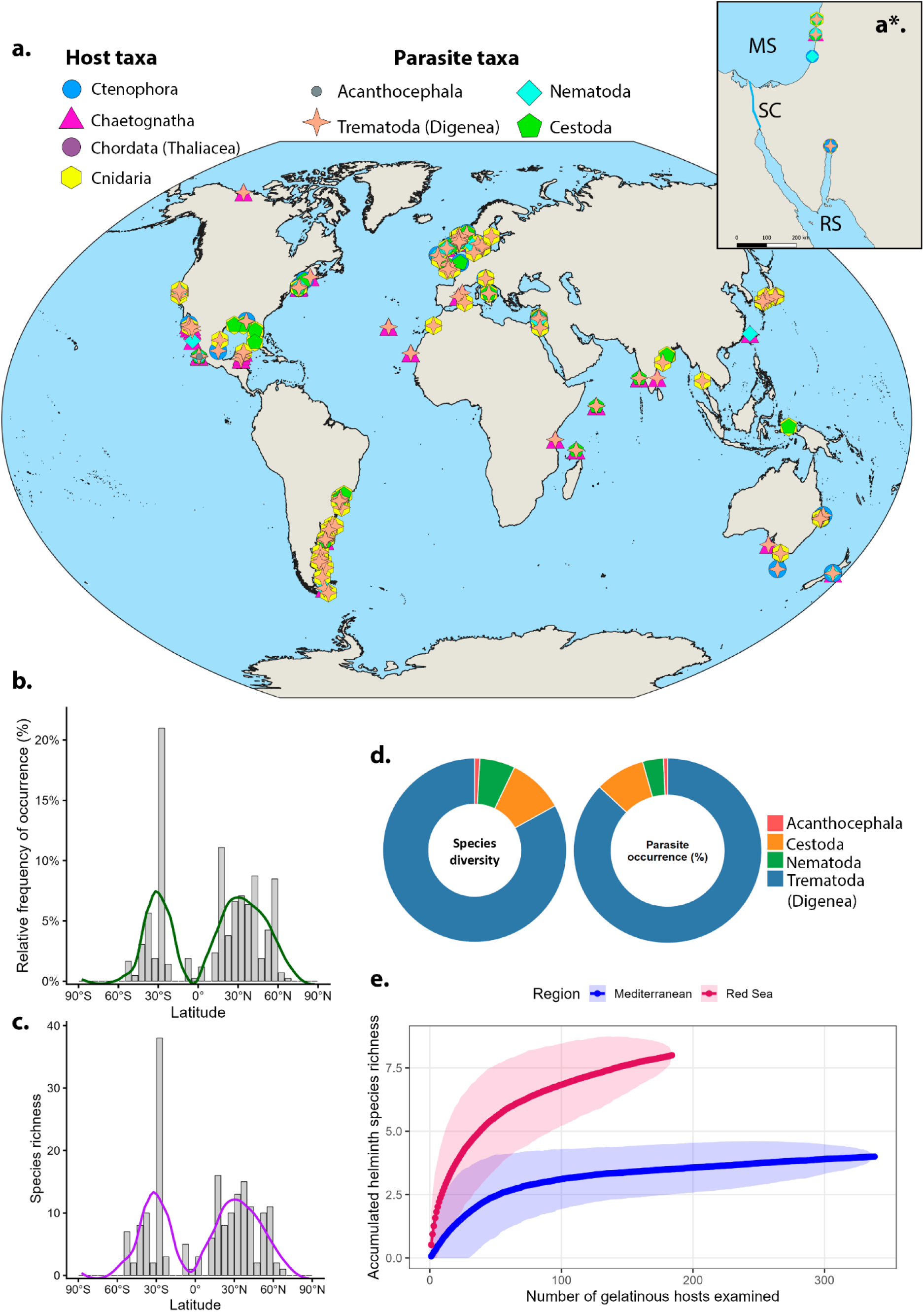
Biogeographic patterns of helminths associated with gelatinous zooplankton. Global (**a**) and regional (**a\***) distribution maps of helminths associated with gelatinous zooplankton. **b** – Relative frequency of helminth occurrences. **c** – Relative species richness. **d** – Helminth composition based on recorded occurrences and diversity. **e** – Species accumulation curve for helminths. For relative frequency of occurrence and species richness, each bin represents 5° of latitude.

Permutation-resampling supported the robustness of the latitudinal pattern. Across 9999 resampled datasets incorporating record-level resampling, latitude jitter, random shifts in bin boundaries and alternative bin widths, occurrence and species-richness peaks remained in temperate latitudes in 96% and 97% of iterations, respectively. Temperate latitudes exceeded tropical latitudes in both occurrence frequency and species richness in all iterations, with median temperate:tropical ratios of 3.79 for occurrence and 3.63 for richness. A complementary null model that retained the number of records per 5° latitude bin but randomized parasite species identities showed that observed richness generally fell within the 95% null intervals expected from record density alone (Supplementary Fig. S1a). The temperate concentration of helminth richness was robust to binning choices and record-level resampling (Supplementary Fig. S1b).

The species-accumulation curves for helminths as a function of the number of gelatinous host individuals examined approached an asymptote at a lower richness level in the Mediterranean Sea than in the Red Sea (Fig. 1e).

### Helminths in gelatinous zooplankton of the Mediterranean Sea

A total of 270 cestode larvae were found parasitizing in 78 *Rhopilema nomadica* hosts from the Mediterranean Sea. All identified cestodes were plerocercoids and occurred exclusively within the host mesoglea (Fig. 2a). They were distributed throughout the bell, oral arms, and, occasionally, the mesoglea of the terminal appendages. Mean body length was 1675.11 ± 595.04 µm, with elongated pale white bodies (Fig. 2b), containing vacuoles, and a scolex armed with small hooks (Fig. 2c, d).

**Figure 2.**
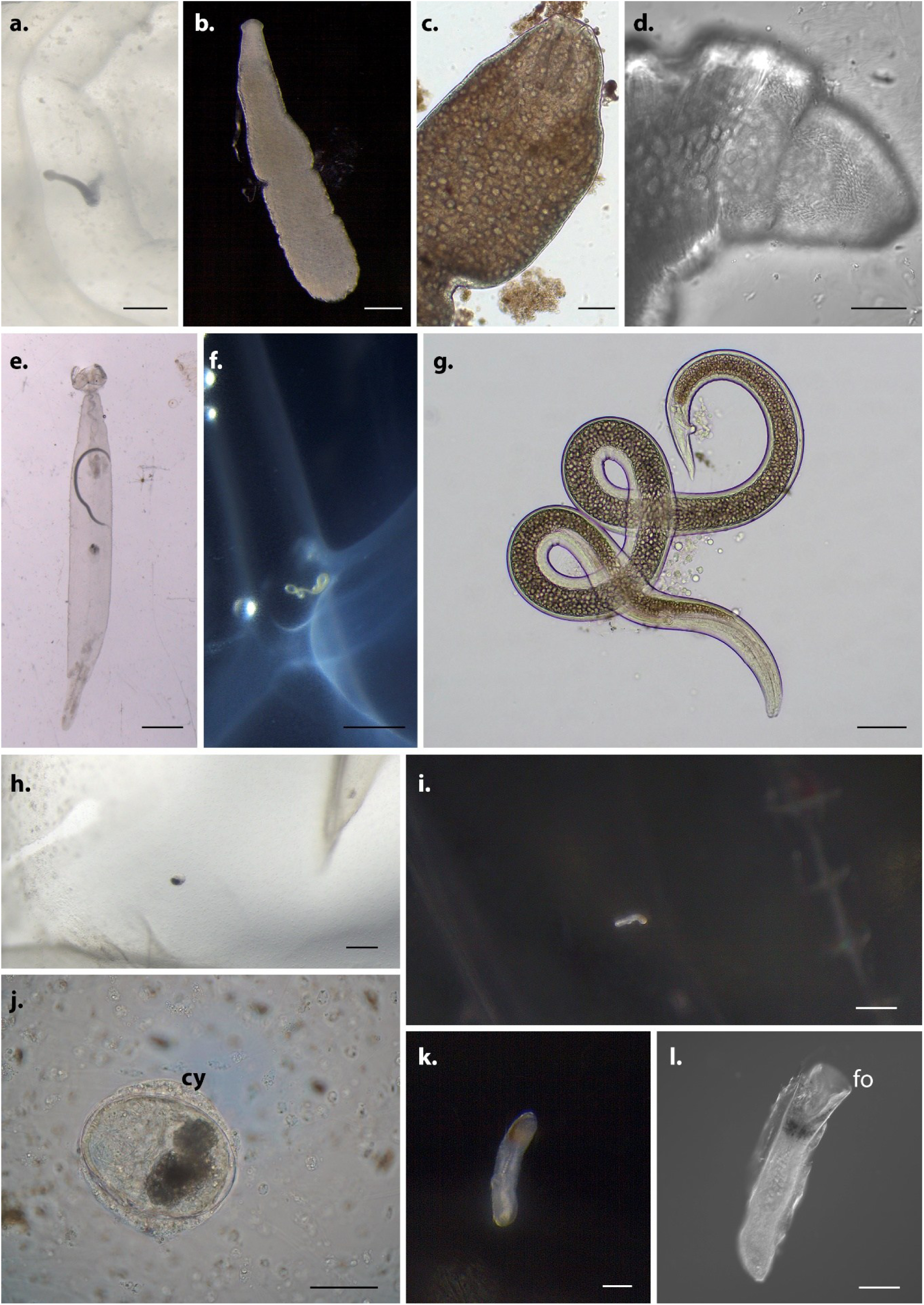
Helminths associated with gelatinous zooplankton from the Mediterranean Sea. **a**-**d** – Cestode *Pseudotobothrium* sp. plerocercoid larvae from *Rhopilema nomadica*. **e**-**g** – Nematode *Hysterothylacium* sp. larvae found in *Flaccisagitta enflata* (**e**), in *R. nomadica* (**f**), and in *Mnemiopsis leidyi* (**g**). **h**, **j** – Encysted metacercariae in mesoglea of *R. nomadica*. cy indicates cyst. **i**, **k**, **l** – Metacercaria of *Opechona kahawai* from *M. leidyi*. fo indicates funnel-shaped oral sucker. Scale bar: a, i – 500 µm, b – 200 µm, c, g, k, l – 100 µm, d, j – 50 µm, e, f – 1000 µm, h – 200 µm.

Twelve nematode larvae were found in *R. nomadica* (5), *Mnemiopsis leidyi* (3), and *Flaccisagitta enflata* (4) (Fig. 2 e-f). In chaetognaths nematodes occurred within the coelom (Fig. 2e), whereas in medusae and ctenophores they were located within the mesoglea (Fig. 2f). Mean body length was 2861.33 ± 548.34 µm.

Twenty-seven trematode metacercariae larvae were found in *R. nomadica* and *M. leidyi*. Most of the metacercariae (26) were encysted within the mesoglea of the bell and oral arms of *R. nomadica* (Fig. 2h, j), with a mean cyst diameter of 69.40 ± 5.98 µm. A single unencysted metacercaria was found in *M. leidyi* (Fig. 2i, k, l). Dark pigmentation in the forebody^14^, together with the broad, funnel-shaped oral sucker (Fig. 2l), originally described by Bray & Cribb^42^ for *Opechona kahawai*, supported its assignment to the family Lepocreadiidae.

### Helminths in gelatinous zooplankton of the Red Sea

Seven gelatinous zooplankton species were found to host metacercariae. The greatest number was found in *Aurelia solida* medusae (1943), followed by the ctenophore *Cestum veneris* (324), the siphonophore *Agalma okenii* (39), the ctenophore *Eurhamphaea vexilligera* (32), the siphonophores *Forskalia tholoides* (28) and *Physophora hydrostatica* (10), and the salp *Thalia* sp. (1). In total, 2377 metacercariae were recorded. Most were unencysted, and some worms from *A. solida* were encysted within the bell mesoglea (Fig. 3a). In siphonophores, metacercariae were located in both nectophores and bracts, whereas in ctenophores, particularly *C. veneris*, most were concentrated along the pharynx (Fig. 3e). We also report, for the first time, a metacercaria within the tunic of the salp *Thalia* sp. (Fig. 3c). Using morphological traits, we assigned the metacercariae to three different families.

**Figure 3.**
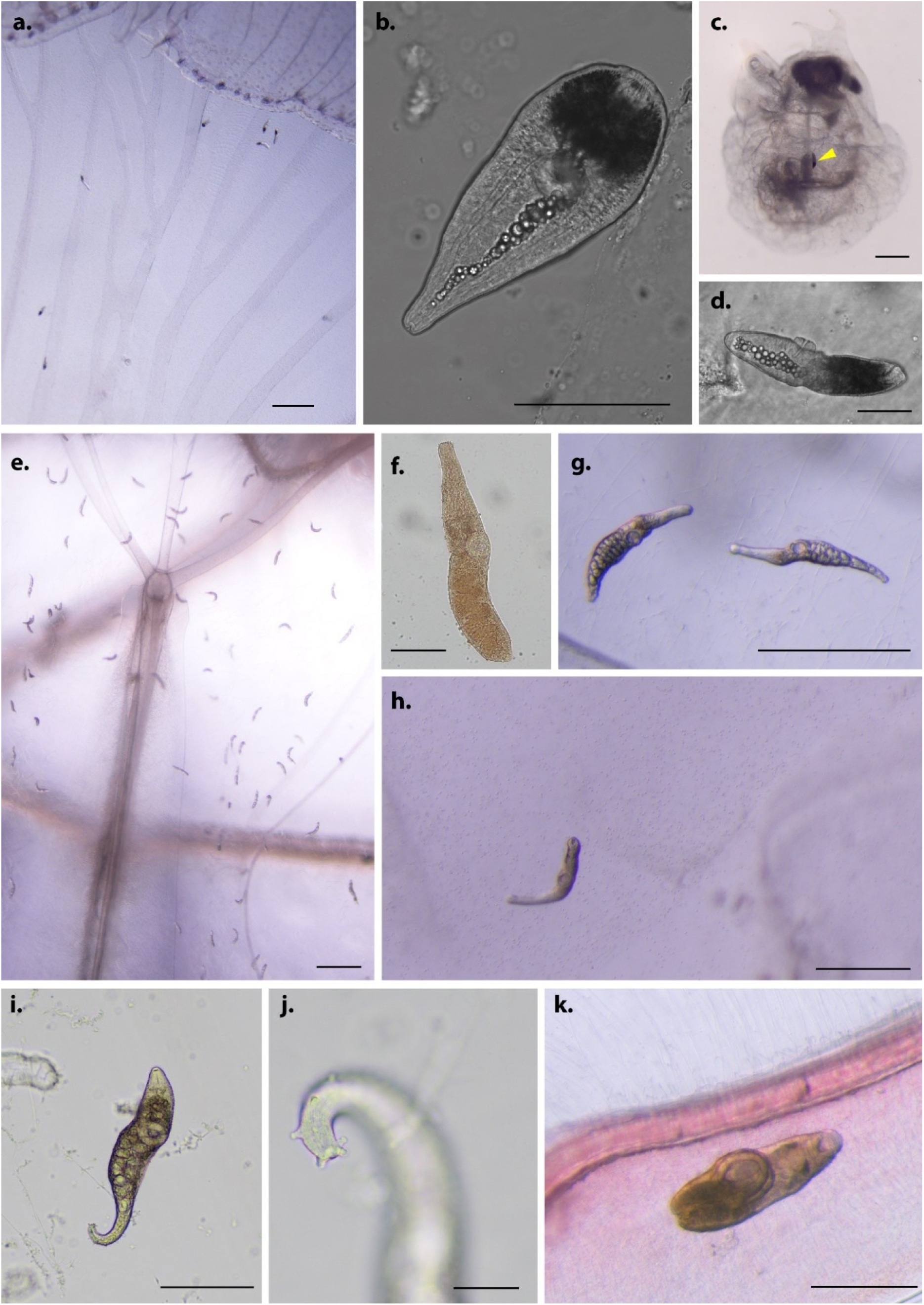
Helminths associated with gelatinous zooplankton from the Red Sea. **a-d** – Metacercariae of *Prodistomum* sp. found in *Aurelia solida* (**a, b, d**) and *Thalia* sp. (yellow arrowhead) (**с**). **e-j** – Didymozoidae metacercariae. *Nematobothrium* sp. in mesoglea of ctenophores *Cestum veneris* (**e, f, g**) and *Eurhamphaea vexilligera* (**i**-**j**). *Didymozoon* sp. in mesoglea of *A. solida* (**h**). **k** – *Odhnerium calyptocotyle* metacercaria in *C. veneris*. Scale bar: **a, e** – 1000 µm, **b, f** – 100 µm, **c, g, h, k** – 500 µm, **d** – 50 µm, **i** – 250 µm, **j** – 25 µm.

The most prevalent and abundant group comprised 2129 lepocreadiid metacercariae (Fig. 3a-d, Supplementary Video 1). These were distinguished from other metacercariae by the dark pigmentation of the anterior part of the body, extending from the oral sucker and almost reaching the ventral sucker (Fig. 3b, d). Both suckers were equal in size; body length was 185.05 ± 55.63 µm. Lepocreadiids also had the broadest host range and were recovered from all seven gelatinous host species.

A second group of metacercariae comprised uncolored, translucent worms with pronounced narrow postacetabular body region and equal-sized suckers (Fig. 3e-j). In total, 248 metacercariae were found in three hosts. Two different morphotypes were distinguished. One occurred in the ctenophores *E. vexilligera* and *C. veneris* (Fig. 3e, f, g, i, j), whereas the other was found exclusively in *A. solida* (Fig. 3h). Metacercariae body size varied between the hosts: in *C. veneris*, worms were 255.78 ± 42.30 µm, in *E. vexilligera* 447.14 ± 49.80 µm, in addition to one exceptionally large specimen of 1065 µm. Metacercariae from *A. solida* had an average size of 264.83 ± 90.12 µm. All metacercariae from ctenophores had papillae-like structures on their bodies, concentrated on the postacetabular part (Fig. 3j, Supplementary Video 2).

A single morphologically-distinct metacercaria was found within the ctenophore *C. veneris*. It measured 717.3 µm in length and had a brown-pigmented body (Fig 3k). Its ventral sucker was approximately 2.5 times larger than the oral sucker. The metacercaria was unencysted and moved freely within the mesoglea (Supplementary Video 3).

### Molecular genetic analyses

#### Cestodes

Analysis of the 28S rRNA gene showed that all the plerocercoids belong to a single species within the genus *Pseudotobothrium* (family Pseudotobothriidae, Trypanorhyncha) (Fig. 4a, Supplementary Fig. S2a). The closest representative is *P. dipsacum* described as plerocercoid in the venus tuskfish *Choerodon venustus* from Australia^43^.

**Figure 4.**
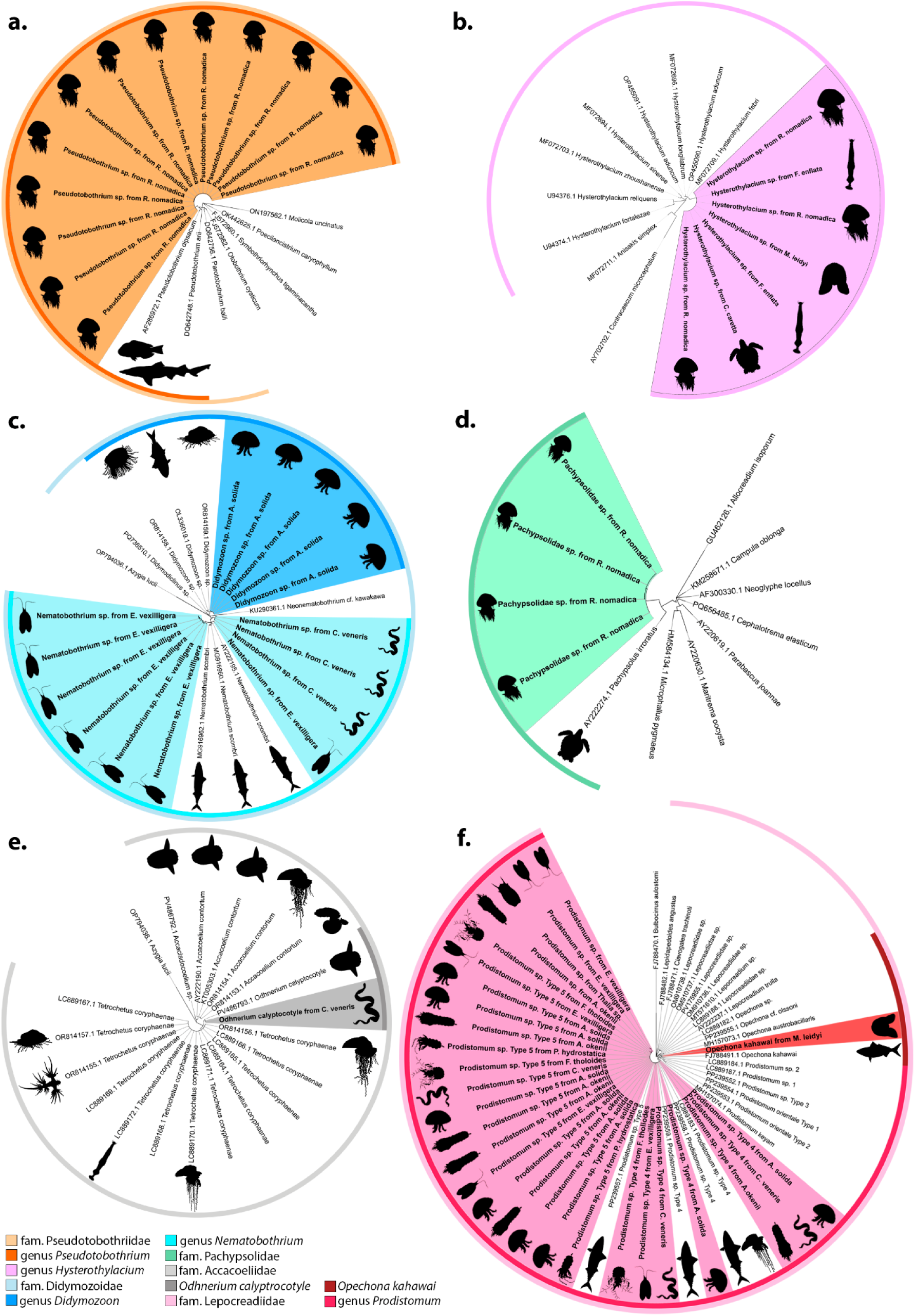
Maximum-likelihood (ML) trees of helminths associated with gelatinous zooplankton. **a** – ML tree of the cestode family Pseudotobothriidae based on 28S rRNA gene. **b** – ML tree of the nematode genus *Hysterothylacium* based on 18S rRNA gene. **c** – ML tree of the trematode family Didymozoidae based on rRNA 28S gene. **d** – ML tree of the trematode family Pachypsolidae based on 28S rRNA gene. **e** – ML tree of the trematode family Accacoeliidae based on 28S rRNA gene. **f** – ML tree of the trematode family Lepocreadiidae based on 28S rRNA gene. Black silhouettes next to helminth names indicate hosts in which they were found. Filled sectors indicate sequences generated in this study.

#### Nematodes

Analysis of 18S rRNA gene indicated that all nematode larvae recovered from gelatinous zooplankton, together with the adult nematodes obtained from the loggerhead sea turtle *Caretta caretta*, were presented by a single species of the genus *Hysterothylacium* (Fig. 4b, Supplementary Fig. S2b). In the maximum likelihood (ML) tree, the nematode sequences obtained in this study formed a strongly supported clade closely related to *H. aduncum* and *H. fabri* (Fig. 4b).

### Digenean trematodes

Analysis of 28S rRNA gene sequences of the specimen obtained from *C. veneris* and other Accacoeliidae trematodes showed a high similarity of metacercaria from this study to *Odhnerium calyptocotyle*. ML tree confirmed that the sequence generated in this study clustered with *O. calyptocotyle* with strong support (Fig. 4e, Supplementary Fig. S2c).

The ML tree based on 28S rRNA gene sequences formed two distinct clades within the family Didymozoidae. The first included metacercariae from *A. solida* and was closely related to *Didymozoon* species previously reported from pleustonic hosts in Australia (Fig. 4c, Supplementary Fig. S3e). The second included sequences from metacercariae recovered from *C. veneris* and *E. vexilligera*, which were placed within the subfamily Nematobothriinae and subdivided into three clusters, related to *Nematobothrium scombri* (Fig. 4c, Supplementary Fig. S3e). Our analysis showed that all lepocreadiid metacercariae from Red Sea hosts form a strongly supported clade with *Prodistomum* sp. types 4 and 5 (Fig. 4f, Supplementary Fig. S3f), originally described by Cribb et al.^14^ in chub mackerel *Scomber japonicus* from Japan and studied by Waki et al.^15^ from gelatinous zooplankton from Japan.

Analysis of 28S rRNA gene of metacercaria found in *M. leidyi* confirmed its identity as *O. kahawai*, showing high similarity to *O. kahawai* described from the Australian salmon kahawai *Arripis trutta* from Tasmanian waters (Fig. 4f, Supplementary Fig. S3f).

Because of the limited availability of related sequences in public databases, the encysted metacercariae recovered from *R. nomadica* could be assigned only to the family Pachypsolidae. Their closest relative was *Pachypsolus irroratus*, described from the olive ridley sea turtle *Lepidochelys olivacea* in Mexico (Fig. 4d, Supplementary Fig. S2d).

### Red-Med parasitological indices

A total of 523 gelatinous zooplankton individuals belonging to 21 species were collected: 184 from the Red Sea and 339 from the Mediterranean Sea. The sampled hosts represented four phyla: Cnidaria (13 species), Ctenophora (5), Chordata (2), and Chaetognatha (1) (Fig. 5). Of these, 109 individuals were found infected by different types of helminths (cestodes, trematodes, and nematodes), including 82 hosts from the Red Sea and 27 from the Mediterranean Sea.

**Figure 5.**
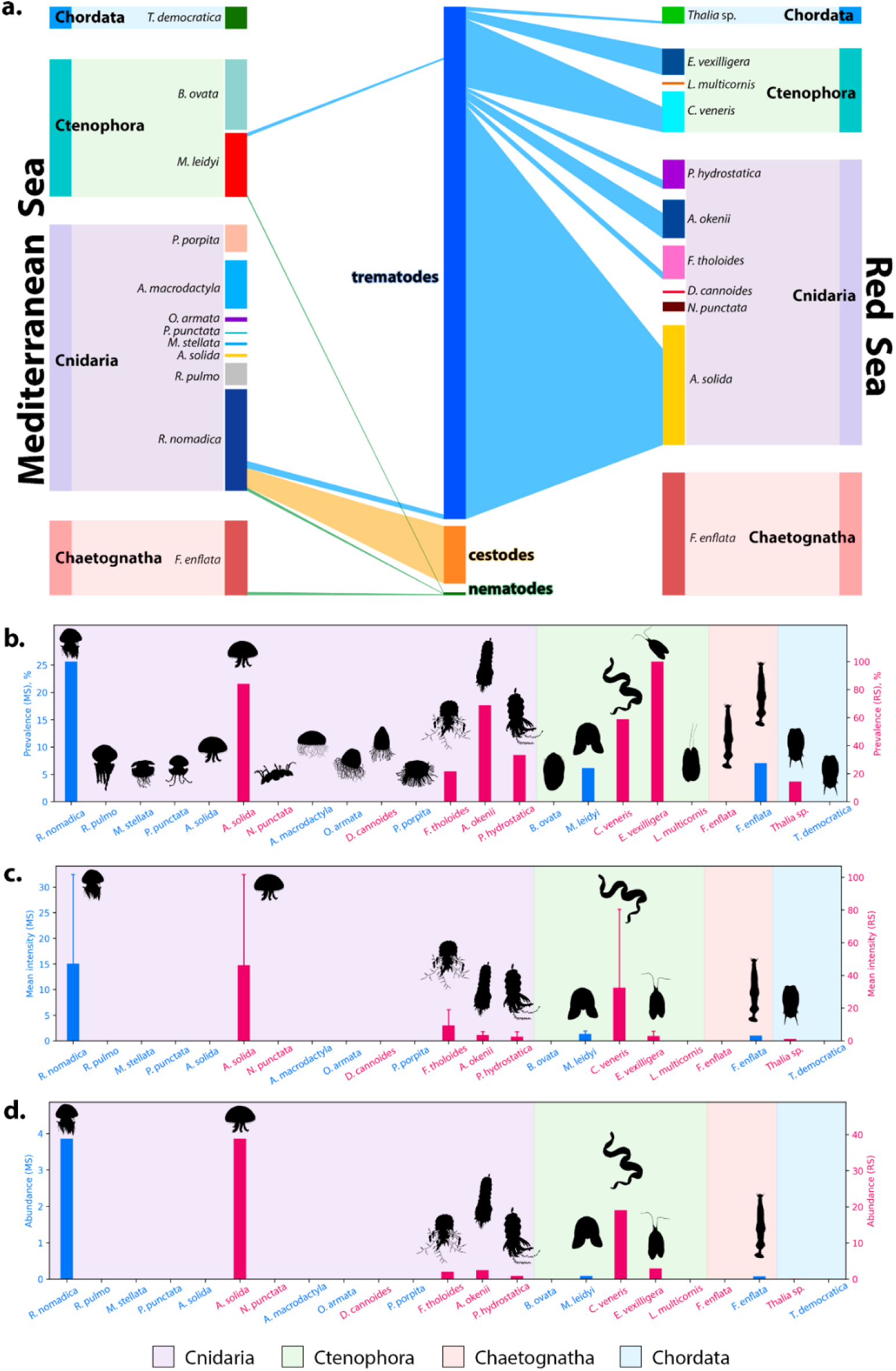
Infection parameters and gelatinous zooplankton host population characteristics. **a** – Bipartite network of host-parasite interactions across two regions – Red and Mediterranean Seas. Host species are positioned on both sides, categorized by region, with parasite species located in the central column. The size of the bars for both hosts and parasites reflect their percentage contribution to overall biodiversity (relative abundance). Connecting lines indicate specific host-parasite associations, with line thickness proportional to the intensity of infection for each host species. Unconnected host species represent taxa in which no parasites were recorded. **b** – Prevalence, mean intensity (**c**), and abundance (**d**) of helminths associated with gelatinous zooplankton. MS – Mediterranean Sea, RS – Red Sea. Host species names in blue indicate gelatinous zooplankton sampled in the Mediterranean Sea, in red – Red Sea.

In total, 2686 helminths were recovered from gelatinous hosts, including 2377 from the Red Sea and 309 from the Mediterranean Sea. Overall parasite prevalence was 44.57% in the Red Sea, compared with 7.99% in the Mediterranean. Mean parasite abundance was 12.92 parasites per host in the Red Sea and 0.91 in the Mediterranean, whereas mean intensity was 28.99 ± 49.40 and 11.44 ± 16.15 parasites per infected host, respectively.

Trematodes dominated the parasite assemblage, accounting for 89.50% of all helminth individuals and exhibiting the greatest taxonomic richness and host breadth (Fig. 5a). Cestodes accounted for 10.05% of all helminths and infected only one host species, whereas nematodes represented 0.45% and infected three host species (Fig. 5a).

Among Red Sea hosts, parasite prevalence was highest in *E. vexilligera* (100%), followed by *A. solida* (84%), *A. okenii* (68.75%), *C. veneris* (58.82%), *P. hydrostatica* (33.33%), *F. tholoides* (21.43%), and *Thalia* sp. (14.29%) (Fig. 5b). In the Mediterranean Sea, prevalence was highest in *R. nomadica* (25.64%), whereas *F. enflata* (7.02%) and *M. leidyi* (6.12%) showed lower rates (Fig. 5b). Mean parasite intensity was highest in *A.* s*olida* (46.26 ± 55.53) and in *C. veneris* (32.40 ± 48.04). Intermediate values were recorded in *R. nomadica* (15.05 ± 17.44) and *F. tholoides* (9.33 ± 9.71), whereas lower intensities were recorded in *A. okenii* (3.55 ± 2.07), *E. vexilligera* (2.91 ± 2.88), *P. hydrostatica* (2.50 ± 3.00), *M. leidyi* (1.33 ± 0.58), *Thalia* sp. (1.00 ± 0), and *F. enflata* (1.00 ± 0) (Fig. 5c.). Mean parasite abundance in the Red Sea was highest in *A. solida* (38.86) and *C. veneris* (19.06). Lower and relatively similar values were recorded in *E. vexilligera* (2.91), *A. okenii* (2.44), and *F. tholoides* (2.00), while *P. hydrostatica* (0.83) and *Thalia* sp. (0.14) had the lowest values (Fig. 5d.). In the Mediterranean, mean abundance was highest in *R. nomadica* (3.86), and considerably lower in *M. leidyi* (0.08) and *F. enflata* (0.07) (Fig. 5d).

No significant relationship between parasite number and host size was detected in the Mediterranean Sea (p>0.05; Supplementary Fig. S4). However, when data were pooled across host species, parasite intensity was significantly related to host size in both the Mediterranean and Red Seas (Supplementary Fig. S4).

## DISCUSSION

Almost all species across the tree of life are involved in parasitic relationships, either as hosts or as parasites, making parasitism one of the most widespread and successful ecological strategies across diverse evolutionary lineages^1,44^. Among this vast diversity, helminths represent one of the most prevalent groups of obligate, highly specialized parasites. Yet, despite their ubiquity, their associations with gelatinous zooplankton have historically received little attention, and the ecological importance of these organisms as helminth hosts has only recently begun to be recognized. Recent studies^17,45^ demonstrated that gelatinous zooplankton might be crucial life-cycle pathway for trematodes infecting pelagic fish. A systematic review^16^ specifically focused on digenean trematodes compiled published records of metacercariae in cnidarians and ctenophores, highlighting their potential as trophic tracers of jellyfish predation and links within marine food webs. Here, we move beyond the group-specific focus of previous studies by providing a global synthesis of all four major helminth groups associated with gelatinous zooplankton, integrating new morphological and molecular observations from the Mediterranean, Red, Celtic, Baltic, and North Seas to reveal broad biogeographic patterns in these associations.

Unlike the classical latitudinal diversity gradient (LDG), which predicts increasing biodiversity toward the equator, numerous terrestrial and aquatic studies have demonstrated that parasite diversity often exhibits an inverse pattern^44,46–49^. The applicability of the LDG remains poorly tested and inadequately explained for parasites within marine ecosystems, where unique environmental and trophic factors may disrupt typical latitudinal trends^50^. Several large-scale analyses have shown that parasite richness and prevalence tend to increase from tropical regions toward temperate latitudes^51,52^, reaching maxima around 30-50° in the Northern Hemisphere. Although these studies focused primarily on helminths recorded from definitive hosts, comparable patterns have also been observed in helminths infecting intermediate hosts^53–55^, suggesting that the phenomenon may represent a general feature of parasite biogeography rather than an artefact of a particular host group or life-cycle stage.

Our global synthesis of helminths associated with gelatinous zooplankton is broadly consistent with this inverse LDG. Parasite records and species richness were higher in temperate regions, whereas tropical latitudes were characterized by a marked scarcity of occurrences. This pattern contrasts with global models of plankton functional groups, which suggested that gelatinous hosts, including jellyfish and salps, generally attain higher richness at low latitudes^56^. It also differs from the global pattern in gelatinous zooplankton biomass^23^, which showed a linear increase from the Southern to the Northern Hemisphere and attributed it to data gaps, the smaller extent of coastline, and lower human influence in the Southern Hemisphere. The inverse LDG observed here is therefore unlikely to simply reflect the diversity or biomass distribution of gelatinous hosts. Instead, the similarity between our results and previously reported large-scale patterns suggests that helminths infecting gelatinous zooplankton follow the same biogeographical processes more characteristic of host–parasite systems than of free-living marine taxa.

A key question, therefore, is whether this pattern reflects a genuine ecological phenomenon or a consequence of incomplete knowledge. The apparent lack of parasite records at low latitudes may partly reflect uneven research intensity, as tropical regions remain comparatively undersampled across a wide range of host taxa, rather than true biological absence^44^. This explanation cannot be dismissed, particularly because both gelatinous hosts and their parasites are notoriously difficult to collect, preserve, and identify. Nevertheless, the repeated observations of similar latitudinal patterns across multiple host taxa, ecosystems, and parasite groups suggest that sampling bias alone is unlikely to explain the observed trend.

One possible mechanism is analogous to the “nasty” host hypothesis^57^. Originally proposed for terrestrial parasitoids, this hypothesis predicts that stronger host defenses reduce the success of specialist consumers. In marine ecosystems, tropical hosts may possess stronger chemical, physiological, or immunological defenses against parasites, potentially decreasing parasite establishment and transmission success. At present, however, empirical evidence supporting this mechanism in marine helminths remains limited, and targeted comparative studies are needed.

A second explanation for the inverse latitudinal gradient in parasites is the resource fragmentation hypothesis^55,58^. This hypothesis proposes that high host diversity in tropical ecosystems leads to the subdivision of available resources among a greater number of host species. As a result, individual host populations tend to be smaller, less abundant, and more spatially fragmented. Because parasites, particularly specialists, rely on host populations as their resource base, fragmented host distributions reduce transmission efficiency and population persistence. Under this framework, lower parasite richness in tropical regions emerges as an indirect consequence of higher host diversity.

Here, we found that the prevalence, intensity and diversity of helminths associated with gelatinous zooplankton were markedly higher in the Red Sea than in the Mediterranean. This contrast suggests that host-parasite interactions may play an important role in the migration dynamics of gelatinous zooplankton in the Mediterranean Sea. One of the most prominent patterns observed in invasive hosts is a reduction in parasite richness and prevalence in the introduced range, consistent with the enemy release hypothesis^32,33^. For Lessepsian migrants, translocation from the Red Sea into the Mediterranean may disrupt parasite transmission pathways, particularly for helminths with complex life cycles requiring multiple intermediate and definitive hosts. Such release from native parasites could provide invasive hosts with fitness advantages by reducing parasite-induced effects on survival, reproduction, or behavior^37^. However, it remains unclear whether the observed pattern represents a true enemy release process or instead reflects broader, more complex biogeographic differences between the two regions that cannot be fully evaluated with our current data. The underlying mechanisms driving these lower infection parameters in the Mediterranean remain open to interpretation and may involve regional variations in environmental conditions, definitive host dynamics, or host community structures.

The ecological consequences of parasite loss may also extend beyond a simple reduction in infection. According to the enemy inversion hypothesis, invasion can reshape species interactions, causing parasites to indirectly benefit introduced hosts^59,60^. For instance, reduced parasite burdens in invasive medusae may improve host condition and bloom potential, enhancing their ecological dominance and expanding transmission pathways through increased biomass availability. Alternatively, parasites acquired in the Mediterranean could integrate these invasive hosts into novel trophic routes, altering regional food web dynamics. Although our data do not directly demonstrate such inversions, the combination of parasite loss and frequent bloom formation suggests that parasite-mediated indirect effects may facilitate invasion success.

Established migrants may also participate in other invasion-related processes. They may acquire native Mediterranean parasites and amplify their transmission to native hosts through spillback, or they may introduce co-invasive parasites that subsequently infect native species through spillover^28^. While our findings primarily support reduced parasite abundance in introduced hosts, these processes cannot be excluded, particularly given the rising dominance of gelatinous zooplankton in the region. As intermediate or paratenic hosts^39^, expanding gelatinous populations remain highly capable of altering existing host-parasite networks across multiple trophic levels.

Marine helminths typically possess complex life cycles requiring transmission through multiple hosts. Gelatinous zooplankton have long been regarded as accidental or dead-end hosts; however, our synthesis of global records together with newly compiled data demonstrates that they represent essential, obligate components of these developmental pathways (Fig. 6). The consistent occurrence of helminths exclusively as larval stages in gelatinous hosts, together with their high prevalence and marked host specificity, indicates that gelatinous zooplankton function as true intermediate hosts rather than merely paratenic vectors. The implications of these findings for the major helminth groups, including cestodes, trematodes and nematodes, are discussed in detail in the Supplementary Discussion.

**Figure 6.**
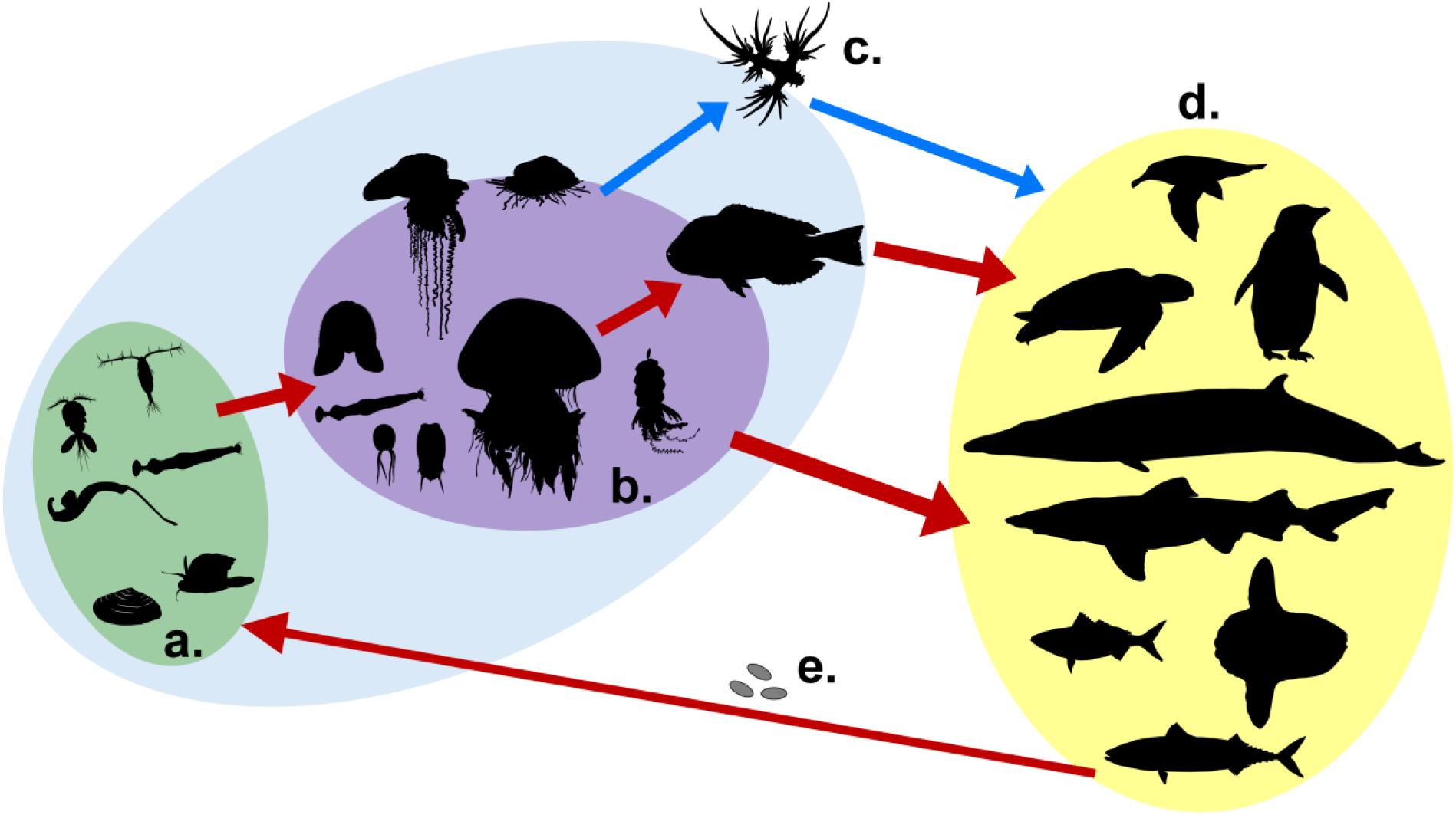
Generalized role of gelatinous zooplankton in helminth transmission pathways. **a** – first benthic or pelagic intermediate host. **b** – second-third (rarely fourth) pelagic intermediate host. **c** – accidental host. **d** – definitive vertebrate host. **e** – helminth eggs released from the definitive host into environment. Red arrows indicate helminth transmission between obligate hosts; blue arrow indicates transmission to/from an accidental host. Green circle indicates first intermediate hosts, purple – second-third intermediate hosts, blue – intermediate hosts in general, where helminth larval stages occur, and yellow – definitive hosts, where helminth adults occur.

More broadly, our study provides a global assessment of helminth diversity, integrating all major helminth groups associated with gelatinous zooplankton, and reveals that these assemblages do not conform to the classical latitudinal diversity gradient typical of many free-living taxa. Instead, they exhibit the inverse latitudinal pattern increasingly reported across host–parasite systems, suggesting that the processes governing parasite diversity differ fundamentally from those shaping free-living biodiversity. Although limited sampling in tropical regions may partly contribute to this pattern, ecological mechanisms such as greater resource fragmentation and stronger host defenses may also play significant roles. Importantly, our study also contributes to addressing one of the major gaps in marine parasitology: more than 95% of marine helminth life cycles remain incompletely known^11^, with different developmental stages often described independently and without reliable molecular links. Resolving these life cycles is essential for identifying transmission pathways, host associations, host switching, and parasite dispersal. By providing molecular data linked to morphologically identified parasite stages and their gelatinous hosts, our study establishes a reference framework that can be used to connect otherwise unrecognized life stages and reconstruct parasite life cycles. Future studies combining standardized sampling across latitudes with quantitative analyses of host community structure and parasite transmission networks will enable us to resolve the relative contributions of these mechanisms to global parasite diversity and to reconstruct the largely unresolved life cycles that underpin these ecological patterns.

## Materials and methods

### Sample collection and examination

Gelatinous zooplankton sampling was performed in May 2020 and between December 2023 and July 2026. Sampling in the Mediterranean Sea was done at the following sites: Hadera (32.4652 °N, 34.8615°E), Olga Beach (32.4389°N, 34.8765°E), Mikhmoret (32.4249°N, 34.8481°E and 32.4023°N, 34.8603°E,) Haifa (32.8264°N, 34.9569°E and 32.8200°N, 34.9537°E), Haifa offshore (32.9310°N, 34.8508 °E), Ashdod (31.8533°N, 34.6431°E). Red Sea sampling point is at the Interuniversity Institute for Marine Sciences (29.5018°N, 34.9179°E) of Eilat. Sampling in the Celtic Sea was done in Cobh (51.849773°N, 8.294108°W). Sampling from the Baltic to the North Sea was performed in Kiel West Shore (54.3234°N, 10.1447°E), Heikendorf (54.3812°N, 10.2007°E), Aalborg (57.0547°N, 9.8724°E), Strandby (57.4961°N, 10.5012°E), and Arendal (58.4602°N, 8.7678°E).

The sampled gelatinous zooplankton comprised the scyphomedusae *Aurelia aurita*, *Aurelia solida*, *Marivagia stellata*, *Nausithoe punctata*, *Phyllorhiza punctata*, *Rhizostoma pulmo*, and *Rhopilema nomadica*; the hydromedusae *Aequorea macrodactyla*, *Aequorea vitrina*, *Dichotomia cannoides*, and *Oceania armata*; the hydrozoan *Porpita porpita*; the siphonophores *Agalma okenii*, *Forskalia tholoides*, and *Physophora hydrostatica*; the ctenophores *Beroe cucumis*, *Beroe ovata*, *Cestum veneris*, *Eurhamphaea vexilligera*, *Leucothea multicornis*, and *Mnemiopsis leidyi*; the chaetognath *Flaccisagitta enflata*; and the salps *Thalia* sp. and *Thalia democratica*.

Medusae, siphonophores, and large ctenophores were collected from the upper 0–3 m using scoop nets, buckets, or ziplock bags to minimize physical damage. Meso- and microzooplankton, including salps, chaetognaths, small ctenophores, and small medusae, were collected using a WP2 net with a 200-µm mesh (Hydro-Bios, Germany). Vertical tows were conducted from a depth of 25 m to the surface, and horizontal tows were conducted over water column depths of 20–70 m by towing at speed 1 knot.

Following collection, specimens were transferred alive to the nearest laboratory for morphometric measurements and parasitological examination. For macrozooplankton, wet mass was recorded, together with bell diameter for medusae, oral–aboral length for ctenophores, and transverse length for *C. veneris*. Siphonophore size could not be precisely inferred due to their extreme fragility. All specimens were examined alive for the presence of helminths. Each individual was first inspected visually. Large medusae and ctenophores were subsequently dissected with a scalpel into sections suitable for examination in Petri dishes. Tissues were inspected using an SZX16 stereomicroscope and a BX43 compound microscope (Olympus, Japan). The presence, number, developmental stage, morphology, and location of helminths within each host were recorded.

Faecal samples from loggerhead (*Caretta caretta*) and green (*Chelonia mydas*) sea turtles were also examined for helminths. Adult nematodes recovered from these samples were used for molecular comparison with nematode larvae obtained from gelatinous zooplankton.

### DNA extraction, amplification and sequencing

Following removal from their hosts, helminths were preserved in absolute ethanol for molecular analysis. Total genomic DNA was extracted using InviSorb Spin Tissue Mini Kit (Invitek Diagnostics, Germany) from whole animals according to the manufacturer’s specifications. The amplifications of the following ribosomal RNA genes were performed: nuclear 18S rRNA gene with primers #3F (5’ GYGGTGCATGGCCGTTSKTRGTT 3’)^61^ and 9R (5’ GATCCTTCCGCAGGTTCACCTAC 3’)^62^ and D1-D3 region of nuclear 28S rRNA gene with primers U178_F (5’ GCACCCGCTGAAYTTAAG 3’) and L1642_R (5’ CCAGCGCCATCCATTTTCA 3’)^63^ for platyhelminths, and nuclear 18S rRNA gene with primer set 18S_98short (5’ GAGGTAGTGACGAAAAATAAC 3’) and 18S_636short (5’ CGTTCTTGATTAATGAAAACATTC 3’)^64^ for nematodes. The nuclear 18S and 28S rRNA genes were selected because their conserved regions permit broad amplification and reliable sequence alignment across diverse helminth lineages, whereas their variable domains provide phylogenetic signal for taxonomic placement. Moreover, both markers are extensively represented in reference databases and have formed the basis of widely accepted phylogenetic frameworks for major helminth groups^65,66^.

Reaction conditions for 18S rRNA gene amplification for platyhelminths (cestodes and trematodes) were as follows: 94 °C for 2 min, followed by 34 cycles of 94 °C for 15 s, 49°C for 30 s, and 72 °C for 1 min, and a final elongation step of 72 °C for 7 min. Reaction conditions for 18S rRNA gene amplification for nematodes: 98 °C for 1 min, followed by 34 cycles of 95 °C for 11 s, 55°C for 10 s, and 72 °C for 10 s, and a final elongation step of 72 °C for 1 min. Reaction conditions for 28S rRNA gene amplification were: 94°C for 2 min followed by 39 cycles of 94°C for 30 s, 52°C for 40 s, and 72°C for 90 s, and a final elongation step of 72°C for 5 min. Obtained PCR products were separated on 1.5% agarose gel and stained with GelRed (Biotium Inc., USA). Purification and Sanger sequencing of the PCR products were performed by Hy Laboratories Ltd. (Rehovot, Israel).

### Bioinformatic and phylogenetic analyses

Sequences were inspected, quality-checked, edited, assembled, and aligned using ClustalW embedded in MEGA v12.0^67^. The best-fitting nucleotide substitution models were selected according to the Bayesian Information Criterion (BIC) using Maximum Likelihood (ML) model selection in MEGA with 1000 bootstrapping replicates to assess branch support. The best-fitting models were K2+G+I for all the platyhelminth datasets and K2+I for the nematode dataset. The sequences generated in this study were deposited in NCBI GenBank under the following accession numbers: nematodes 18S (PZ745763-PZ745769), platyhelminths 18S (PZ745936-PZ745945, PZ750763), platyhelminths 28S (PZ750701-PZ750762). ML trees were visualised using Interactive Tree of Life tool (iTOL v6)^68^.

#### Cestodes

A total of twenty 28S rRNA gene sequences of Trypanoselachoida were analysed, including thirteen sequences of *Pseudotobothrium* sp. plerocercoids obtained in this study and seven from GenBank. A sequence of *Molicola uncinatus* (ON197562) was used as outgroup.

#### Digenean trematodes

##### Accacoeliidae

A total of twenty 28S rRNA gene sequences of Hemiurata were analysed, including one sequence of *Odhnerium calyptocotyle* obtained in this study and nineteen from GenBank. A sequence of *Azygia lucii* (OP794036) was used as outgroup.

##### Didymozoidae

The dataset of twenty-four Hemiurata 28S rRNA gene sequences, including fifteen sequences obtained in this study and nine from GenBank, was analyzed. A sequence of *Azygia lucii* (OP794036) was used as outgroup.

##### Lepocreadiidae

Sixty-one 28S rRNA gene sequences of Lepocreadioidea were analysed: twenty-nine obtained in this study and thirty-two from GenBank. A sequence of *Bulbocirrus aulostomi* (FJ788470) was used as outgroup.

##### Pachypsolidae

Twelve 28S rRNA gene sequences of Xiphidiata were analysed: four obtained in this study and eight from GenBank. A sequence of *Allocreadium isoporum* (GU462126) was used as outgroup.

#### Nematodes

A total of seventeen 18S rRNA gene sequences of Ascaridoidea were analysed, including seven *Hysterothylacium* sequences obtained in this study and ten from GenBank. A sequence of *Contracaecum microcephalum* (AY702702) was used as outgroup.

### Global systematic review

A systematic literature search for records of helminths associated with gelatinous zooplankton was conducted in the ISI Web of Science and Scopus databases on 9 April 2026. The search combined keywords describing gelatinous hosts with terms describing helminth parasites: (“gelatinous zooplankton* OR jellyfish* OR ctenophore* OR comb-jell* OR combjell* OR chaetognath* OR siphonophore* OR medusa* OR salp* OR thalia* OR doliolid*) AND (helminth* OR trematode* OR digenea* OR nematode* OR cestode* OR acanthocephala*”). Publications in English, French, German, Portuguese, Italian, and Russian were considered.

Following removal of duplicate records, titles and abstracts were screened for relevance, and potentially eligible publications were evaluated using the full text. Additional publications not recovered by the database search were identified through backward and forward citation tracking of relevant papers. Studies were included when they reported: (1) a gelatinous zooplankton host, (2) a helminth parasite, and (3) an association originating from a marine or brackish-water environment. Studies involving non-gelatinous hosts, non-helminth symbionts, or exclusively freshwater or terrestrial environments were excluded. Taxonomic names of both hosts and helminth parasites were verified against the World Register of Marine Species (WoRMS) and standardized to currently accepted nomenclature; unresolved identifications and taxa of uncertain status were retained.

For each eligible record, we extracted the publication and sampling year; study location and geographic coordinates; gelatinous host taxon; parasite taxon and developmental stage; number of hosts examined and infected; number of parasites recorded; prevalence; mean abundance; and mean intensity. When coordinates were not reported directly, they were assigned based on the most precise locality information available in the publication. Collected data and data generated in this study were combined to construct a global distribution map with helminth-infected gelatinous zooplankton records using QGIS Geographic Information System^69^.

We calculated the relative reported occurrence of each helminth taxonomic group as the proportion of all parasite-association records assigned to that group. To characterize latitudinal patterns, records were assigned to 5° latitude bins. The relative frequency of records within each bin was calculated as the proportion of all records occurring in that bin. Parasite species richness was calculated as the number of unique parasite taxa recorded within each 5° bin. Parasite species occurring in multiple latitudinal bins were counted once in each bin in which they occurred.

To evaluate the robustness of the observed latitudinal patterns, we performed a permutation-resampling sensitivity analysis. The analysis was repeated across 9999 resampled datasets. In each iteration, occurrence records were resampled with replacement, latitudes were randomly jittered by ±2.5°, bin boundaries were randomly shifted, and latitudinal patterns were recalculated using alternative bin widths of 5°, 10° and 15°. For each iteration, we recorded the latitude of maximum occurrence frequency and maximum species richness, and calculated the ratio of temperate to tropical occurrence frequency and species richness. Tropical latitudes were defined as |latitude| < 23.5°, and temperate latitudes as 23.5°≤ |latitude| ≤ 60°.

In addition, we tested whether the observed latitudinal richness pattern could be explained solely by uneven numbers of occurrence records among latitude bins. For this null model, parasite species identities were randomly permuted among records while retaining the observed latitude and the number of records per latitude bin. Observed helminth species richness in each bin was then compared with the null distribution generated from 9999 permutations, and standardized effect sizes were calculated.

### Parasitological indices and species accumulation curves

For each host species and region, parasite prevalence was calculated as the percentage of examined hosts that were infected. Mean abundance was calculated as the total number of parasites divided by the total number of examined hosts, including uninfected individuals. Mean intensity was calculated as the total number of parasites divided by the number of infected hosts.

Parasite species accumulation curves were generated separately for Mediterranean and Red Sea host samples to compare the rate at which parasite richness increased with sampling effort. Species accumulation curves were calculated using random permutations of host order with the specaccum function in the vegan package in R^70,71^. Curves were generated independently for each region, with 999 permutations used to estimate mean accumulated parasite richness and confidence intervals.

## Data availability

All sequence data generated in this work were deposited in the National Center for Biotechnology Information (NCBI GenNank) under accession numbers PZ745763-PZ745769 (nematodes 18S), PZ745936-PZ745945 and PZ750763 (platyhelminths 18S), PZ750701-PZ750762 (platyhelminths 28S) and will be made publicly available upon publication. The systematic review dataset is fully detailed in Supplementary Tables S1 and S2. R scripts used for data processing and analyses are available at https://github.com/guy-haimlab/helminths_in_gz.

## Acknowledgements

A. Iakovleva acknowledges support from the Bloom Scholarship for Doctoral Studies from the Bloom Graduate School, University of Haifa; the Mediterranean Sea Research Center of Israel (tender 24/3); and a 2024–2025 research grant from the Interuniversity Institute for Marine Sciences in Eilat (IUI). We thank the National Monitoring Program of the Israel Oceanographic and Limnological Research, the Israel Nature and Parks Authority for providing sampling permit no. 2025/43831, and the Sea Turtle Rescue Center in Mikhmoret for providing samples and assistance with this research. We are grateful to the members of the Guy-Haim Zooplankton Ecology Lab at IOLR and to A.R. Morov for their help. We also thank R. Kiko, M. Wahl, and T. Reusch for providing access to laboratory facilities at GEOMAR Helmholtz Centre for Ocean Research, and the IOLR and IUI sea-going teams for their assistance with sampling. We thank B. Sures, F.R. Barboza, and M.J. Costello for helpful discussions about the results and their interpretation. A. Iakovleva expresses her deepest gratitude to N. Pismennyi for the endless support throughout her life and at every stage of the research.

## Supplementary Discussion

Our findings on cestodes associated with gelatinous zooplankton constitute one of the most extensive datasets reported to date, whereas previous records have been sparse, geographically fragmented, and rarely synthesized. This knowledge gap may partly reflect linguistic barriers, because many early European observations were published in local-language sources and have remained relatively inaccessible to the broader research community.

The biogeography of the cestodes identified here suggests an Indo-Pacific affinity. Larvae of *Pseudotobothrium dipsacum* have previously been reported from coral reef-associated fish in Australia^1,2^. A decade later, the same larvae have been identified in the teleost fish from the southern Red Sea near the Bab-el-Mandeb Strait, where the basin connects with the Indian Ocean^3^. Adult cestodes of the same genus, *P. arii*, have likewise been recorded in the broadfin shark *Lamiopsis temminckii* from Malaysia^4^.

Notably, these cestodes were detected exclusively in *Rhopilema nomadica* among all gelatinous hosts examined, suggesting a relatively narrow host association at this life-history stage. This pattern is particularly intriguing because *R. nomadica* is an invasive species of Indo-Pacific origin^5^. It is highly plausible that these medusae co-introduced the parasites during their introduction to the Mediterranean Sea. Furthermore, our data sheds light on the life history of the family Pseudotobothriidae^6^. The developmental pathway of these cestodes appears to involve three intermediate hosts – a planktonic crustacean, a gelatinous zooplankter, and a teleost fish. Within the gelatinous zooplankton acting as the second intermediate host, the larva continues its development, transforming into an elongated plerocercoid stage localized in the host mesoglea. The evolutionary choice of this specific host category depends on the cestode species and the dietary preferences of the subsequent hosts in the trophic chain^7^. Upon ingestion of medusa by the teleost fish, and eventually by elasmobranch definitive hosts, the plerocercoid matures into an adult tapeworm, which aligns with the typical life-cycle patterns of the order Trypanorhyncha^8^.

Nematodes of the genus *Hysterothylacium* exhibit a cosmopolitan distribution, utilizing a remarkably wide array of marine hosts^9,10^. Their larvae have previously been recorded mainly from chaetognaths, ctenophores^11^, and hydromedusae^9^. In the present study, they were detected across a diverse range of gelatinous hosts, further confirming their exceptionally low host specificity. Notably, this includes the first documentation of *Hysterothylacium* sp. in the scyphomedusa *R. nomadica*, a finding that further underscores the high plasticity of their life-cycle pathways. Furthermore, the occurrence of the same nematodes in Mediterranean sea turtles implies that these reptiles regularly include gelatinous zooplankton in their diet^12^, confirming a viable and predictable transmission route for these generalist parasites within the regional pelagic food web.

In contrast to cestodes and nematodes, digenean trematodes represent the most extensively studied and frequently documented group of parasites associated with gelatinous zooplankton. Their complex life cycles typically require sequential transmission through mollusc first intermediate hosts, planktonic second intermediate hosts, and various vertebrate definitive hosts. Within our datasets, this species-rich group was represented by the families Lepocreadiidae, Accacoeliidae, Didymozoidae, and Pachypsolidae.

Lepocreadiidae was the most prevalent and broadly distributed trematode family found in gelatinous hosts. While Cribb et al.^13^ documented only adult stages of the closely related *Prodistomum* types 4 and 5 in their definitive host, the chub mackerel *S. japonicus* in Japan, we discovered their metacercariae across a wide range of gelatinous hosts, infecting nearly all seven gelatinous zooplankton species examined in the Red Sea. This widespread occurrence highlights low host specificity of these larvae within the region. Although *S. japonicus* is not found in the Red Sea, a closely related congener, the blue mackerel *Scomber australasicus*, is well-documented in the region^14^. This geographic overlap suggests a potential life-cycle pathway where *S. australasicus* might substitute *S. japonicus* as the local definitive host, facilitating the transmission of these trematodes. This alternative route is further supported by Ansai et al.^15^ and Waki et al.^16^, who traced the same *Prodistomum* type 4 sporocysts in the gastropod *Purpuradusta gracilis* to metacercariae infecting diverse Pacific hosts, such as *Chrysaora pacifica*, *Physalia utriculus*, and *Urashimea globosa*. Interestingly, *S. australasicus* may also provide a biogeographic link for *Opechona kahawai*. This trematode was originally described from Australian waters, where it infects the endemic fish *Arripis trutta*^17^, the yellowtail amberjack *Seriola lalandi*^18^, and *S. australasicus*^19^. Because *S. australasicus* is distributed throughout the Indo-Pacific, including the Red Sea, it may represent a potential definitive host linking the native range of *O. kahawai* with the Mediterranean. Although this hypothesis requires molecular confirmation, the occurrence of *O. kahawai* in the Mediterranean suggests that the parasite could have dispersed through the Indo-Pacific–Red Sea corridor by exploiting different definitive hosts. Collectively, these observations support an Indo-Pacific origin for these trematodes and indicate that mackerels may serve as important definitive hosts. Browne et al.^19^ hypothesized that gelatinous zooplankton are critical for the successful completion of the life cycles of lepocreadiid trematodes. Our results provide substantial evidence supporting this hypothesis. Notably, the discovery of these metacercariae in the salp *Thalia* sp. not only expands the known potential pathways of transmission but also represents the first record of helminths within pelagic tunicates globally.

Accumulating evidence suggests that members of the family Accacoeliidae consistently utilize gelatinous zooplankton as essential intermediate hosts to successfully reach their definitive hosts. Adults of this family are well-known obligate parasites of ocean sunfishes (family Molidae)^20^, and other large pelagic predators, such as mahi-mahi *Coryphaena hippurus*. In particular, adult stages of *Odhnerium calyptocotyle* have been documented during parasitological surveys of *Mola mola* in the Sea of Okhotsk and the Sea of Japan^20,21^. We report the first documentation of *O. calyptocotyle* metacercaria within the ctenophore *C. veneris*, contributing to our understanding of this helminth’s life cycle. Although our single record limits broad conclusions, combining this finding with existing records of *Tetrochetus coryphaenae* and *Accacoelium contortum* metacercariae in pleustonic and pelagic cnidarians^16,22^ allows us to infer that accacoeliids rely on gelatinous zooplankton as obligate vectors. Given that definitive hosts like ocean sunfishes are known consumers of gelatinous zooplankton^23^, our data suggest that they serve as the primary transmission vector that effectively bridges the trophic gap to these apex pelagic predators, a hypothesis similarly proposed by Louvard et al.^22^.

Members of the digenean trematode family Didymozoidae are widely distributed among marine teleost fishes, yet their life cycles remain largely unresolved. Didymozoids constitute the most morphologically diverse group among trematodes^24^. Historically, Nikolaeva^25^ proposed that molluscs and planktonic invertebrates act as the first and second intermediate hosts, respectively. Decades later, this hypothetical cycle has received empirical support. Regarding the first intermediate hosts, Louvard et al.^26^ confirmed that didymozoid transmission is driven by the trophic ecology of a mollusс host. The pelagic pathway involving crustaceans as the second host, and gelatinous zooplankton as third or subsequent intermediate hosts was demonstrated by records in pleuston from Australia^22^ and from the Gulf of California^27,28^. The high numbers of metacercariae found in pleustonic cnidarians strongly suggested that these hosts are critical links in the pelagic trophic transmission pathway^22^. Didymozoid larvae consistently exhibited both high prevalence and intensity, alongside repetitive seasonal occurrences and a high degree of host specificity, further confirming this hypothesis and supporting the role of gelatinous zooplankton as obligate intermediate hosts in didymozoid life cycles.

Since its original description, the family Pachypsolidae has remained a monophyletic yet enigmatic taxon with entirely unknown life cycles^29^. While adults are known to infect caimans and sea turtles – specifically documented in the loggerhead *Caretta caretta* from the Mediterranean Sea and the olive ridley *Lepidochelys olivacea* from Mexico^29,30^ – their developmental pathways remained unresolved. Here, we present the first record of pachypsolid metacercariae parasitizing an intermediate planktonic host. Our identification of new representatives of this family utilizing the gelatinous host *R. nomadica*, with sea turtles acting as the plausible definitive hosts including medusae in their diets^12^, provides critical insights that finally sheds light on the life cycle of this family. This finding suggests a high host specificity at the intermediate stage, indicating that these parasites may selectively utilize specific gelatinous taxa to optimize transmission to their definitive hosts.

---

**Supplementary Figure S1.**
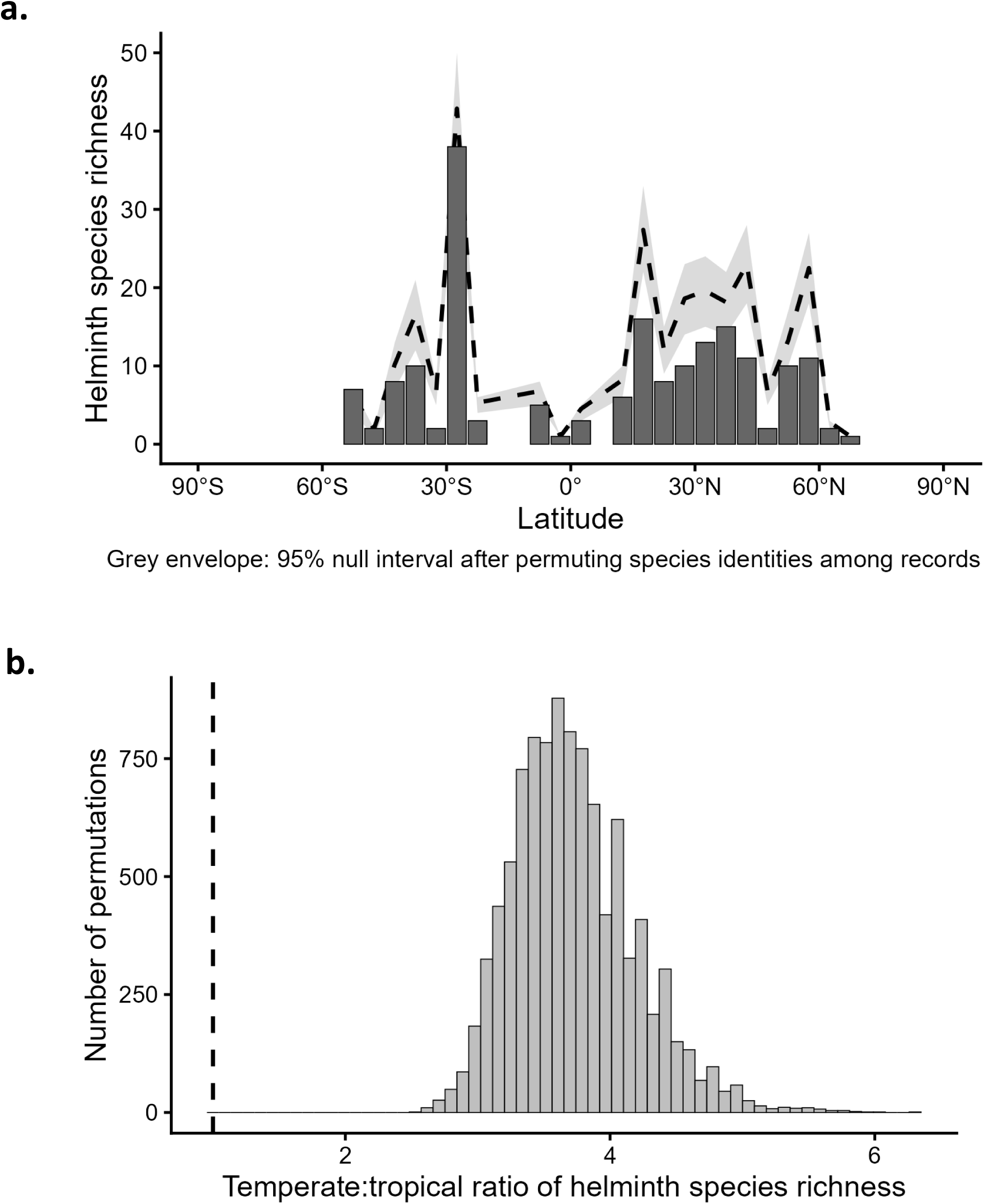
Sensitivity analyses of the latitudinal diversity pattern. **a –** Null model of helminth species richness across 5° latitudinal bins based on 9999 permutations. Bars show observed richness, the dashed line shows mean permuted richness, and the grey envelope indicates the 95% null interval. **b –** Permutation-resampling distribution of the temperate to tropical species richness ratio. Values greater than 1 indicate higher helminth richness in temperate than tropical latitudes. The dashed line marks equal richness.

**Supplementary Figure S2.**
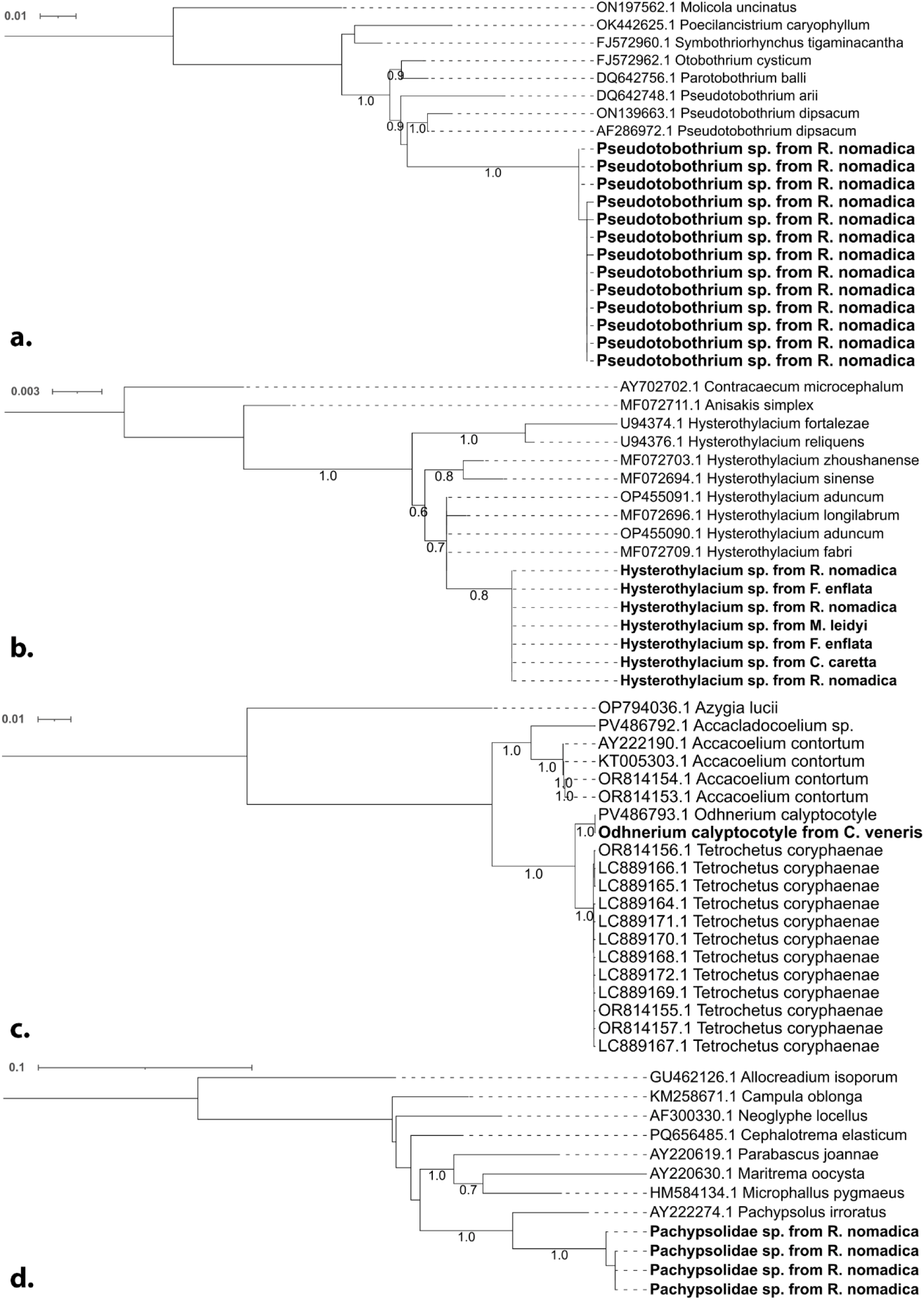
Maximum-likelihood (ML) trees of helminths associated with gelatinous zooplankton. **a** – ML tree of the cestode family Pseudotobothriidae based on 28S rRNA gene. **b** – ML tree of the nematode genus *Hysterothylacium* based on 18S rRNA gene. **c** – ML tree of the trematode family Accacoeliidae based on 28S rRNA gene. **d** – ML tree of the trematode family Pachypsolidae based on 28S rRNA gene. The numbers below the branches indicate the proportion of ML bootstrap support (1000 replicates) for nodes that received at least 60% support. Names in bold indicate sequences generated in this study.

**Supplementary Figure S3.**
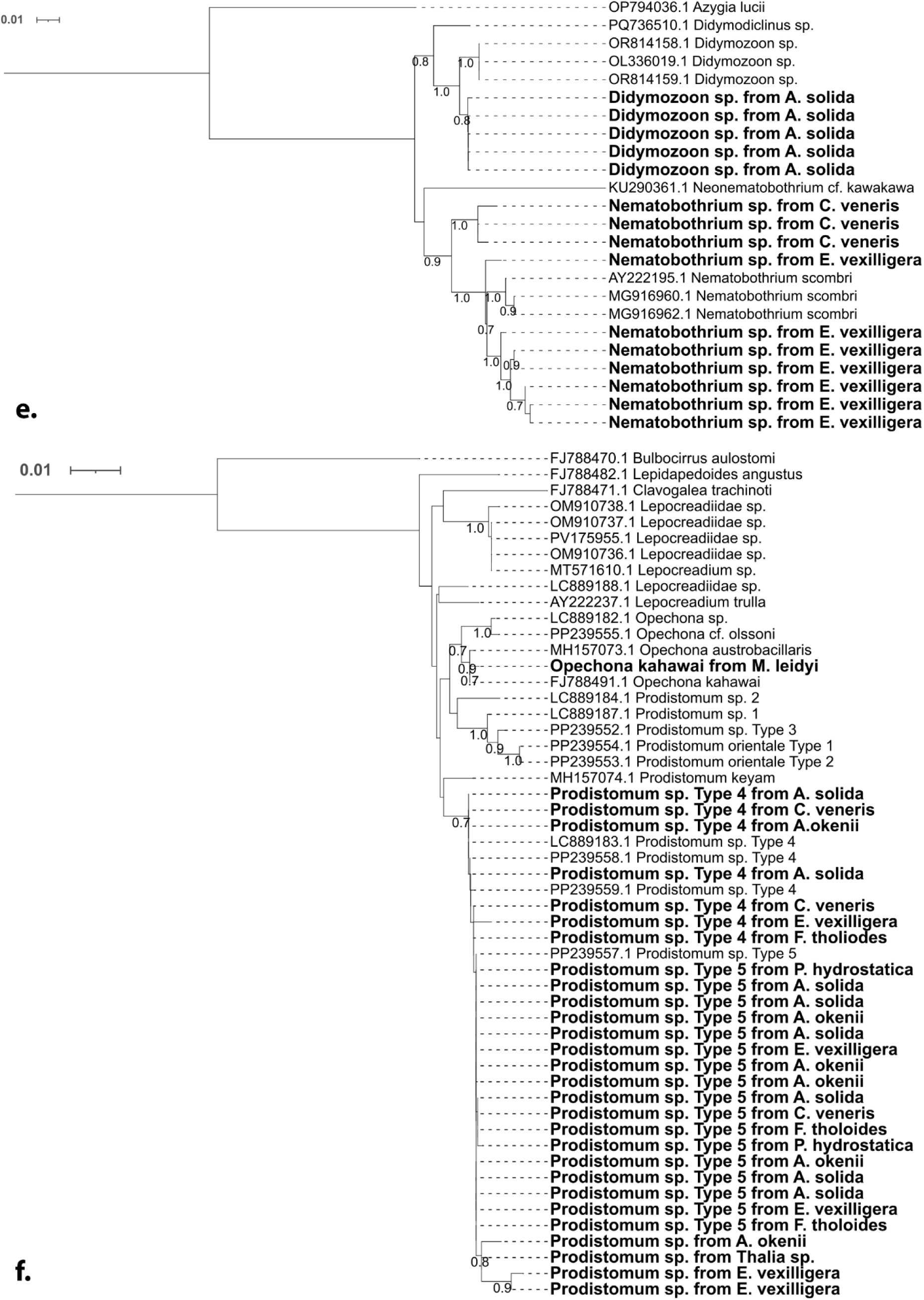
Maximum-likelihood (ML) trees of helminths associated with gelatinous zooplankton. **e** – ML tree of the trematode family Didymozoidae based on 28S rRNA gene. **f** – ML tree of the trematode family Lepocreadiidae based on 28S rRNA gene. The numbers below the branches indicate the proportion of ML bootstrap support (1000 replicates) for nodes that received at least 60% support. Names in bold indicate sequences generated in this study.

**Supplementary Figure S4.**
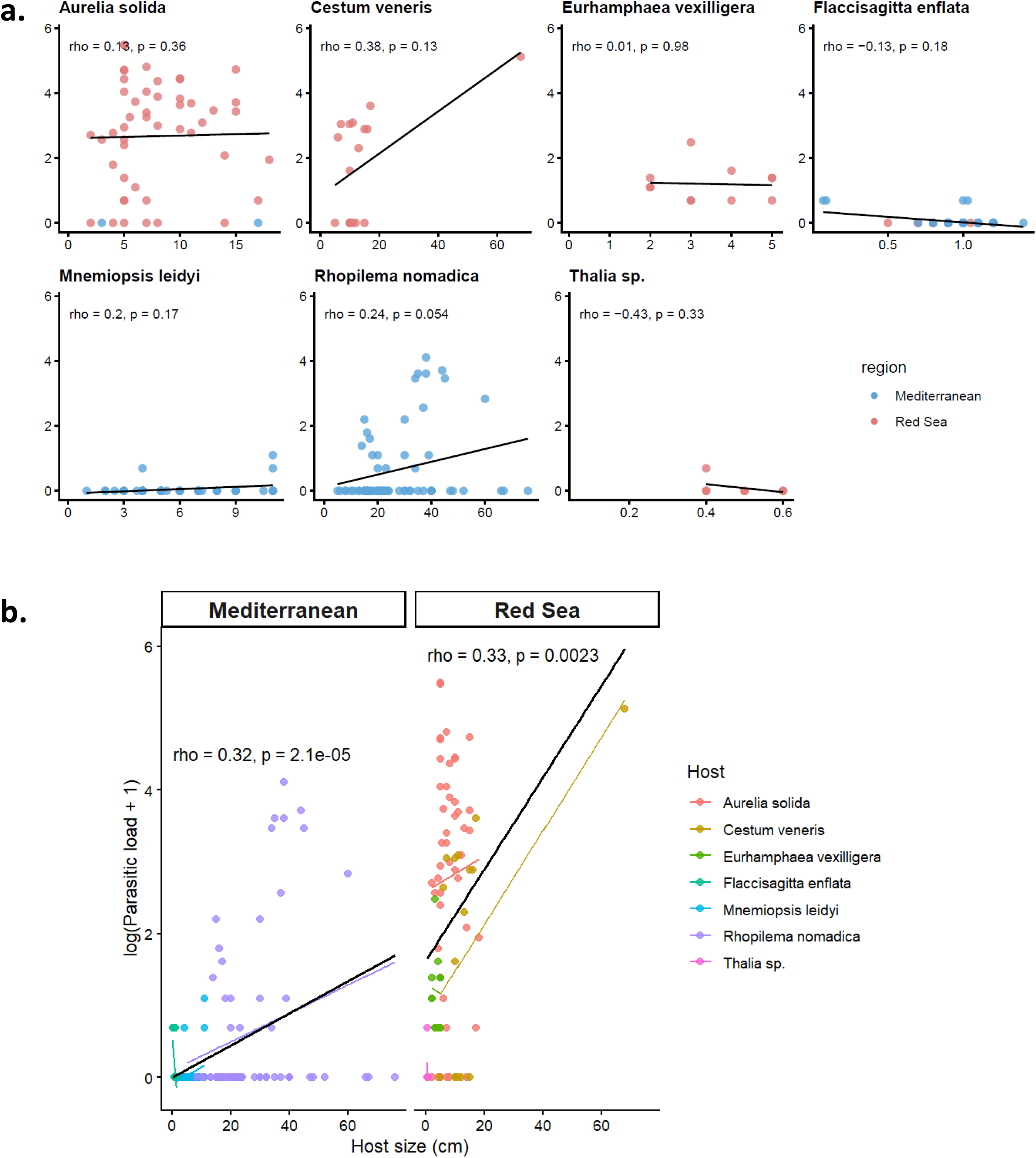
Correlation (Spearman’s rank) between gelatinous host size (cm) and the number of parasites (infection intensity) per host species (log+1) (**a**) and per region (**b**).

**Supplementary Table S1.** Documented 431 records of helminth associations with gelatinous zooplankton hosts extracted from global systematic review and this study.

**Supplementary Table S2.** Global systematic review dataset showing the list of 89 retained references (1830-2026). Only relevant references are shown.

**Supplementary video 1.** *Prodistomum* sp. in *Aurelia solida* host. Red Sea, March 2024.

**Supplementary video 2.** *Nematobothrium* sp. in *Eurhamphaea vexilligera* host. Red Sea, September 2025.

**Supplementary video 3.** *Odhnerium calyptocotyle* in *Cestum veneris* host. Red Sea, March 2025.

## References

1 Poulin, R. & Morand, S. The Diversity of Parasites. The Quarterly Review of Biology 75 (2000). 10.1086/393500

2 Poulin, R. The functional importance of parasites in animal communities: many roles at many levels? International journal for parasitology 29, 903–914 (1999). 10.1016/s0020-7519(99)00045-4

3 Byers, J. E. Marine Parasites and Disease in the Era of Global Climate Change. Annual Review of Marine Science 13 (2021). 10.1146/annurev-marine-031920-100429

4 Smit, N. J. & Sures, B. Aquatic Parasitology: Ecological and Environmental Concepts and Implications of Marine and Freshwater Parasites. 1 edn, (Springer Cham, 2025).

5 Dobson, A., Lafferty, K. D., Kuris, A. M., Hechinger, R. F. & Jetz, W. Homage to Linnaeus: How many parasites? How many hosts? Proceedings of the National Academy of Sciences 105 (2008). 10.1073/pnas.0803232105

6 Thomas, F., Poulin, R., de Meeüs, T., Guégan, J.-F. & Renaud, F. Parasites and ecosystem engineering: what roles could they play? Oikos, 167–171 (1999). 10.2307/3546879

7 Arias-González, J. E. & Morand, S. Trophic functioning with parasites: a new insight for ecosystem analysis. Marine Ecology Progress Series 320 (2006). 10.3354/meps320043

8 Dunne, J. A. et al. Parasites Affect Food Web Structure Primarily through Increased Diversity and Complexity. PLOS Biology 11 (2013). 10.1371/journal.pbio.1001579

9 Lafferty, K. D. et al. Parasites in food webs: the ultimate missing links. Ecology Letters 11 (2008). 10.1111/j.1461-0248.2008.01174.x

10 Sures, B., Díaz-Morales, D. M., Yong, R. Q.-Y., Erasmus, A. & Schwelm, J. Biology and Life Cycle of Helminths. Aquatic Parasitology: Ecological and Environmental Concepts and Implications of Marine and Freshwater Parasites (2025). 10.1007/978-3-031-83903-0_5

11 Blasco-Costa, I. & Poulin, R. Parasite life-cycle studies: a plea to resurrect an old parasitological tradition. Journal of Helminthology 91 (2017). 10.1017/S0022149X16000924

12 Nogueira Júnior, M., et al. Diversity of gelatinous zooplankton (Cnidaria, Ctenophora, Chaetognatha and Tunicata) from a subtropical estuarine system, southeast Brazil. Marine Biodiversity 2018 49:3 49 (2018). 10.1007/s12526-018-0912-7

13 Lozano-Cobo, H. et al. Finding a needle in a haystack: larval stages of Didymozoidae (Trematoda: Digenea) parasitizing marine zooplankton. Parasitology Research 2022 121:9 121 (2022). 10.1007/s00436-022-07593-6

14 Cribb, T. H. et al. Lepocreadiidae (Trematoda) associated with gelatinous zooplankton (Cnidaria and Ctenophora) and fishes in Australian and Japanese waters. Parasitology International 101 (2024). 10.1016/j.parint.2024.102890

15 Waki, T. et al. Metacercariae infecting seven cnidarian species with their life cycle information including their adult stages in Japan. Journal of Helminthology 100 (2026). 10.1017/S0022149X25100989

16 Pitt, K. A. et al. Digenean parasites as novel tracers of predation on jellyfish. Marine Environmental Research 213 (2026). 10.1016/j.marenvres.2025.107647

17 Browne, J. G., Pitt, K. A. & Cribb, T. H. DNA sequencing demonstrates the importance of jellyfish in life cycles of lepocreadiid trematodes. Journal of Helminthology 94 (2020). 10.1017/S0022149X20000632

18 Ohtsuka, S. et al. Symbionts of marine medusae and ctenophores. Plankton and Benthos Research 4 (2009). 10.3800/pbr.4.1

19 Ohtsuka, S. et al. In-situ observations of symbionts on medusae occurring in Japan, Thailand, Indonesia and Malaysia. 広島大学総合博物館研究報告, 9–18 (2010).

20 Kondo, Y. et al. Seasonal changes in infection with trematode species utilizing jellyfish as hosts: evidence of transmission to definitive host fish via medusivory. Parasite 23, 16 (2016). 10.1051/parasite/2016016

21 Artsdatabanken. Metazoan parasites of non-crustacean zooplankton project (ParaZoo) (The Norwegian Biodiversity Data Bank, University Museum of Bergen, University of Bergen, 2022–2024).

22 Lebrato, M., Molinero, J.-C., Mychek-Londer, J. G., Gonzalez, E. M. & Jones, D. O. B. Gelatinous Carbon Impacts Benthic Megafaunal Communities in a Continental Margin. Frontiers in Marine Science 9 (2022). 10.3389/fmars.2022.902674

23 Lucas, C. H. et al. Gelatinous zooplankton biomass in the global oceans: geographic variation and environmental drivers. Global Ecology and Biogeography 23 (2014). 10.1111/geb.12169

24 Luo, J. Y. et al. Gelatinous Zooplankton-Mediated Carbon Flows in the Global Oceans: A Data-Driven Modeling Study. Global Biogeochemical Cycles 34 (2020). 10.1029/2020GB006704

25 Guy-Haim, T. et al. The effects of decomposing invasive jellyfish on biogeochemical fluxes and microbial dynamics in an ultra-oligotrophic sea. Biogeosciences 17 (2020). 10.5194/bg-17-5489-2020

26 Hays, G. C., Doyle, T. K. & Houghton, J. D. R. A Paradigm Shift in the Trophic Importance of Jellyfish? Trends in Ecology & Evolution 33 (2018). 10.1016/j.tree.2018.09.001

27 Sousa, W. P. & Grosholz, E. D. The influence of habitat structure on the transmission of parasites. Habitat Structure (1991). 10.1007/978-94-011-3076-9_15

28 Iakovleva, A. et al. From ctenophores to scyphozoans: parasitic spillover of a burrowing sea anemone. Scientific Reports 2024 14:1 14 (2024). 10.1038/s41598-024-72168-7

29 Dunn, A. M. Chapter 7 Parasites and Biological Invasions. Advances in Parasitology 68 (2009). 10.1016/S0065-308X(08)00607-6

30 Dunn, A. M. et al. Indirect effects of parasites in invasions. Functional Ecology 26 (2012). 10.1111/j.1365-2435.2012.02041.x

31 Dunn, A. M. & Hatcher, M. J. Parasites and biological invasions: parallels, interactions, and control. Trends in Parasitology 31 (2015). 10.1016/j.pt.2014.12.003

32 Torchin, M. E. et al. Introduced species and their missing parasites. Nature 2003 421:6923 421 (2003). 10.1038/nature01346

33 Chalkowski, K., Lepczyk, C. A. & Zohdy, S. Parasite Ecology of Invasive Species: Conceptual Framework and New Hypotheses. Trends in Parasitology 34 (2018). 10.1016/j.pt.2018.05.008

34 Lymbery, A. J., Morine, M., Kanani, H. G., Beatty, S. J. & Morgan, D. L. Co-invaders: The effects of alien parasites on native hosts. International Journal for Parasitology: Parasites and Wildlife 3 (2014). 10.1016/j.ijppaw.2014.04.002

35 Díaz-Morales, D. M., Sures, B., Jolma, E. R. & Thieltges, D. W. Invasion Biology in the Context of Aquatic Host–Parasite Interac. Aquatic Parasitology: Ecological and Environmental Concepts and Implications of Marine and Freshwater Parasites (2025). 10.1007/978-3-031-83903-0_18

36 Galil, B. S. A Sea Under Siege – Alien Species in the Mediterranean. Biological Invasions 2 (2000). 10.1023/A:1010057010476

37 Diamant, A., Goren, M., Galil, B., Yokes, B. & Klopman, Y. Parasites of Red-Med immigrant and native Mediterranean coastal fish species: new observations from the Israeli and Turkish coasts. Rapp Comm Int Mer Medit 39, 495 (2010).

38 Edelist, D. et al. Phenological shift in swarming patterns of *Rhopilema nomadica* in the Eastern Mediterranean Sea. Journal of Plankton Research 42 (2020). 10.1093/plankt/fbaa008

39 Marcogliese, D. J. The role of zooplankton in the transmission of helminth parasites to fish. Reviews in Fish Biology and Fisheries 1995 5:3 5 (1995). 10.1007/BF00043006

40 Marcogliese, D. J. Food webs and the transmission of parasites to marine fish. Parasitology 124 (2002). 10.1017/S003118200200149X

41 Thiel, M. E. Wirbellose Meerestiere als Parasiten, Kommensalen oder Symbionten in oder an Scyphomedusen. Helgoländer wissenschaftliche Meeresuntersuchungen 1976 28:3 28 (1976). 10.1007/BF01610591

42 Bray, R. & Cribb, T. New species of *Opechona* Looss, 1907 and *Cephalolepidapedon* Yamaguti, 1970 (Digenea: Lepocreadiidae) from fishes off northern Tasmania. Papers and Proceedings of the Royal Society of Tasmania (2003). 10.26749/rstpp.137.1

43 Olson, P., Littlewood, D., Bray, R. & Mariaux, J. Interrelationships and Evolution of the Tapeworms (Platyhelminthes: Cestoda). Molecular Phylogenetics and Evolution 19 (2001). 10.1006/mpev.2001.0930

44 Poulin, R. & Leung, T. L. F. Latitudinal gradient in the taxonomic composition of parasite communities. Journal of Helminthology 85 (2011). 10.1017/S0022149X10000696

45 Louvard, C., Yong, R. Q.-Y., Cutmore, S. C. & Cribb, T. H. The oceanic pleuston community as a potentially crucial life-cycle pathway for pelagic fish-infecting parasitic worms. International Journal for Parasitology 54 (2024). 10.1016/j.ijpara.2023.11.001

46 Choudhury, A. & Dick, T. A. Richness and diversity of helminth communities in tropical freshwater fishes: empirical evidence. Journal of Biogeography 27 (2000). 10.1046/j.1365-2699.2000.00450.x

47 Poulin, R. Another look at the richness of helminth communities in tropical freshwater fish. Journal of Biogeography 28 (2001). 10.1046/j.1365-2699.2001.00570.x

48 Krasnov, B. R., Shenbrot, G. I., Khokhlova, I. S. & Degen, A. A. Flea species richness and parameters of host body, host geography and host ‘milieu’. Journal of Animal Ecology 73 (2004). 10.1111/j.0021-8790.2004.00883.x

49 Johnson, P. & Haas, S. E. Why do parasites exhibit reverse latitudinal diversity gradients? Testing the roles of host diversity, habitat and climate. Global Ecology and Biogeography 30 (2021). 10.1111/geb.13347

50 Poulin, R. & Thieltges, D. W. Patterns and Processes in Marine Parasite Biogeography and Macroecology. The Ecology and Evolution of Marine Parasites and Disease, 125 (2026). 10.1093/9780197790847.003.0008

51 Poulin, R. Latitudinal gradients in parasite diversity: bridging the gap between temperate and tropical areas. Neotropical Helminthology 4, 169–177 (2010).

52 Morris, T. C., Costello, M. J. & Matzke, N. J. Marine Parasite Biogeography Mirrors Host Patterns Across Latitude, Area, and Diversity. New Zealand Journal of Zoology 53 (2026). 10.1002/njz2.70032

53 Thieltges, D. W., Fredensborg, B. L., Studer, A. & Poulin, R. Large-scale patterns in trematode richness and infection levels in marine crustacean hosts. Marine Ecology Progress Series 389 (2009). 10.3354/meps08188

54 Moore, C., McCoy, K. & McCoy, M. Latitudinal Variation in Trematode Prevalence Across Regions in North and South America: Evidence of an Inverse Gradient in Second-Intermediate and Final Hosts. Journal of Biogeography 53 (2026). 10.1111/jbi.70157

55 Torchin, M. E., Miura, O. & Hechinger, R. F. Parasite species richness and intensity of interspecific interactions increase with latitude in two wide-ranging hosts. Ecology 96 (2015). 10.1890/15-0518.1

56 Benedetti, F., Gruber, N. & Vogt, M. Global gradients in species richness of marine plankton functional groups. Journal of Plankton Research 45 (2023). 10.1093/plankt/fbad044

57 Gauld, I. D., Gaston, K. J. & Janzen, D. H. Plant allelochemicals, tritrophic interactions and the anomalous diversity of tropical parasitoids: the “nasty” host hypothesis. Oikos, 353–357 (1992). 10.2307/3545032

58 Kindlmann, P., Schödelbauerová, I. & Dixon, A. Inverse latitudinal gradients in species diversity. Scaling Biodiversity, 246 – 257 (2007). 10.1017/CBO9780511814938.014

59 Pearson, D. E. & Callaway, R. M. Indirect effects of host-specific biological control agents. Trends in Ecology & Evolution 18 (2003). 10.1016/S0169-5347(03)00188-5

60 Colautti, R. I., Ricciardi, A., Grigorovich, I. A. & MacIsaac, H. J. Is invasion success explained by the enemy release hypothesis? Ecology Letters 7 (2004). 10.1111/j.1461-0248.2004.00616.x

61 Machida, R. J. & Knowlton, N. PCR Primers for Metazoan Nuclear 18S and 28S Ribosomal DNA Sequences. PLOS ONE 7 (2012). 10.1371/journal.pone.0046180

62 Giribet, G., Carranza, S., Baguñà, J., Riutort, M. & Ribera, C. First molecular evidence for the existence of a Tardigrada+ Arthropoda clade. Molecular biology and evolution 13, 76–84 (1996). 10.1093/oxfordjournals.molbev.a025573

63 Lockyer, A. E. et al. The phylogeny of the Schistosomatidae based on three genes with emphasis on the interrelationships of *Schistosoma* Weinland, 1858. Parasitology 126 (2003). 10.1017/S0031182002002792

64 Harbuzov, Z. et al. Amplicon sequence variant-based meiofaunal community composition revealed by DADA2 tool is compatible with species composition. Marine Genomics 65 (2022). 10.1016/j.margen.2022.100980

65 Olson, P., Cribb, T., Tkach, V., Bray, R. & Littlewood, D. Phylogeny and classification of the Digenea (Platyhelminthes: Trematoda). International journal for parasitology 33, 733–755 (2003).

66 Chan, A. H. E., Chaisiri, K., Saralamba, S., Morand, S. & Thaenkham, U. Assessing the suitability of mitochondrial and nuclear DNA genetic markers for molecular systematics and species identification of helminths. Parasites & Vectors 2021 14:1 14 (2021). 10.1186/s13071-021-04737-y

67 Kumar, S. et al. MEGA12: Molecular Evolutionary Genetic Analysis Version 12 for Adaptive and Green Computing. Mol Biol Evol 41 (2024). 10.1093/molbev/msae263

68 Letunic, I. & Bork, P. Interactive Tree of Life (iTOL) v6: recent updates to the phylogenetic tree display and annotation tool. Nucleic Acids Research 52 (2024). 10.1093/nar/gkae268

69 QGIS Geographic Information System v. 3.34.6 (QGIS Association, 2024).

70 R Core Team R: a language and environment for statistical computing (2025).

71 vegan: Community ecology package (Version 2.7-2)[Software] (2025).

## Supplementary Discussion References

1 Beveridge, I., Bray, R. A., Cribb, T. H. & Justine, J.-L. Diversity of trypanorhynch metacestodes in teleost fishes from coral reefs off eastern Australia and New Caledonia. Parasite 21 (2014 Nov 18). 10.1051/parasite/2014060

2 Olson, P., Littlewood, D., Bray, R. & Mariaux, J. Interrelationships and Evolution of the Tapeworms (Platyhelminthes: Cestoda). Molecular Phylogenetics and Evolution 19 (2001). 10.1006/mpev.2001.0930

3 Al-Zubaidy, A. B. & Mhaisen, F. T. Larval tapeworms (Cestoda: Trypanorhyncha) from some Red Sea fishes, Yemen. Mesopotamian Journal of Marine Sciences 26 (2011). 10.58629/mjms.v26i1.186

4 Olson, P. D. et al. Evolution of the trypanorhynch tapeworms: Parasite phylogeny supports independent lineages of sharks and rays. International Journal for Parasitology 40 (2010/02/01). 10.1016/j.ijpara.2009.07.012

5 Kuplik, Z., et al. *Rhopilema nomadica* in the Mediterranean: Molecular Evidence for Migration and Insights into Its Proliferation. Diversity *2026, Vol. 18*, Page 94 18 (2026). 10.3390/d18020094

6 Palm, H., Waeschenbach, A., Olson, P. & Littlewood, D. Molecular phylogeny and evolution of the Trypanorhyncha (Platyhelminthes: Cestoda). Molecular Phylogenetics and Evolution 52 (2009). 10.1016/j.ympev.2009.01.019

7 Sures, B., Díaz-Morales, D. M., Yong, R. Q.-Y., Erasmus, A. & Schwelm, J. Biology and Life Cycle of Helminths. Aquatic Parasitology: Ecological and Environmental Concepts and Implications of Marine and Freshwater Parasites (2025). 10.1007/978-3-031-83903-0_5

8 Palm, H. in Proceedings of the IVth international workshop on theoretical and marine parasitology, AtlantNIRO, Kaliningrad. 164–170.

9 Marcogliese, D. J. The role of zooplankton in the transmission of helminth parasites to fish. Reviews in Fish Biology and Fisheries 1995 5:3 5 (1995). 10.1007/BF00043006

10 Tedesco, P. et al. *Hysterothylacium fabri* (Nematoda: Raphidascarididae) in *Mullus surmuletus* (Perciformes: Mullidae) and *Uranoscopus scaber* (Perciformes: Uranoscopidae) from the Mediterranean. The Journal of Parasitology 104 (2018). 10.1645/17-115

11 Køie, M. Aspects of the life cycle and morphology of *Hysterothylacium aduncum* (Rudolphi, 1802) (Nematoda, Ascaridoidea, Anisakidae). Canadian Journal of Zoology 71 (1993). 10.1139/z93-178

12 Arai, M. N. Predation on pelagic coelenterates: a review. Journal of the Marine Biological Association of the United Kingdom 85 (2005). 10.1017/S0025315405011458

13 Cribb, T. H. et al. Lepocreadiidae (Trematoda) associated with gelatinous zooplankton (Cnidaria and Ctenophora) and fishes in Australian and Japanese waters. Parasitology International 101 (2024). 10.1016/j.parint.2024.102890

14 Baker, E. A., Collette, B. B., Baker, E. A. & Collette, B. B. Mackerel from the Northern Indian Ocean and the Red Sea are *Scomber australasicus*, not *Scomber japonicus*. Ichthyological Research 1998 45:1 45 (1998). 10.1007/BF02678572

15 Ansai, E. et al. Life cycles of trematodes infecting six species of intertidal gastropods in Japan. Journal of Helminthology 99 (2025). 10.1017/S0022149X25100485

16 Waki, T. et al. Metacercariae infecting seven cnidarian species with their life cycle information including their adult stages in Japan. Journal of Helminthology 100 (2026). 10.1017/S0022149X25100989

17 Bray, R. & Cribb, T. New species of *Opechona* Looss, 1907 and *Cephalolepidapedon* Yamaguti, 1970 (Digenea: Lepocreadiidae) from fishes off northern Tasmania. Papers and Proceedings of the Royal Society of Tasmania (2003). 10.26749/rstpp.137.1

18 Hutson, K., Ernst, I., Mooney, A. & Whittington, I. Metazoan parasite assemblages of wild *Seriola lalandi* (Carangidae) from eastern and southern Australia - PubMed. Parasitology international 56 (2007). 10.1016/j.parint.2006.12.003

19 Browne, J. G., Pitt, K. A. & Cribb, T. H. DNA sequencing demonstrates the importance of jellyfish in life cycles of lepocreadiid trematodes. Journal of Helminthology 94 (2020). 10.1017/S0022149X20000632

20 Waki, T., Nakano, H., Okawa, T. & Ishikawa, T. Accacoeliid trematodes in two sunfish species, *Masturus lanceolatus* and *Mola mola*, in Japanese waters. Systematic Parasitology 103 (2026). 10.1007/s11230-025-10259-3

21 Sokolov, S. G., Gordeev, I. I. & Atopkin, D. M. Phylogeny of two accacoeliid species (Digenea: Hemiuroidea) ex *Mola mola* (Linnaeus, 1758) (Tetraodontiformes: Molidae) from Northwest Pacific, with first molecular data on *Odhnerium* Yamaguti, 1934. Journal of Helminthology 99 (2025). 10.1017/S0022149X25000380

22 Louvard, C., Yong, R. Q.-Y., Cutmore, S. C. & Cribb, T. H. The oceanic pleuston community as a potentially crucial life-cycle pathway for pelagic fish-infecting parasitic worms. International Journal for Parasitology 54 (2024). 10.1016/j.ijpara.2023.11.001

23 Chang, C.-T. et al. Diet breadth and overlap in the Family Molidae. Environmental Biology of Fishes 107 (2024). 10.1007/s10641-024-01582-7

24 Meneses, Y. C. et al. Integrative Taxonomy of Didymozoids Parasitizing *Thunnus obesus* (Scombridae) from Southwest Atlantic Ocean: A New Genus and Species. Pathogens 2025, Vol. 14, Page 359 14 (2025). 10.3390/pathogens14040359

25 Nikolaeva, V. On the development cycle of trematodes belonging to the family Didymozoidae (Monticelli, 1888) Poche, 1907. Zoologicheski Zhurnal 44, 1317–1327 (1965).

26 Louvard, C., Corner, R. D., Cutmore, S. C. & Cribb, T. H. Evidence that host ecology drives first intermediate host use in the Didymozoidae (Trematoda: Hemiuroidea): an asexual infection in a vermetid (Gastropoda). Journal of Helminthology 96 (2022). 10.1017/S0022149X22000748

27 Lozano-Cobo, H., Gutiérrez, J. G. & Robinson, C. J. Didymozoid trematode parasitizing *Pleurobrachia bachei* (Ctenophora) and *Muggiaea atlantica* (Siphonophora) in the Gulf of California: Didymozoid in gelatinous zooplankton. Hidrobiológica 35 (2025).

28 Lozano-Cobo, H. et al. Finding a needle in a haystack: larval stages of Didymozoidae (Trematoda: Digenea) parasitizing marine zooplankton. Parasitology Research 2022 121:9 121 (2022). 10.1007/s00436-022-07593-6

29 Olson, P., Cribb, T., Tkach, V., Bray, R. & Littlewood, D. Phylogeny and classification of the Digenea (Platyhelminthes: Trematoda). International journal for parasitology 33, 733–755 (2003).

30 Marchiori, E. et al. An extensive survey on helminth community of *Caretta caretta* from the neritic feeding grounds of Northwestern Adriatic sea. Scientific Reports 2025 15:1 15 (2025). 10.1038/s41598-025-15272-6

